# Deciphering the patterns and timing of diversification of the genus *Melanastera* (Hemiptera: Psylloidea: Liviidae) in the Neotropics

**DOI:** 10.1101/2024.12.30.630774

**Authors:** Liliya Štarhová Serbina, Daniel Burckhardt, Lenka Petráková Dušátková, Dalva L. Queiroz, Renato Goldenberg, Hannes Schuler, Diana M. Percy, Igor Malenovský

## Abstract

Even after decades of research on diversification in the Neotropics, our understanding of the evolutionary processes shaping Neotropical clades is still incomplete. In the current study, we used different divergence times and likelihood-based methods to investigate the influence of biogeography and host associations on the diversification of the most species-rich Neotropical psyllid genus *Melanastera* (Liviidae) using molecular phylogenetic data from seven gene fragments (four mitochondrial and three nuclear). The putatively monophyletic group of Neotropical *Melanastera* species has an estimated crown node age of 20.2 Ma (ML, CI 20.2–30.6) or 23.2 Ma (BI, 95% HPD 16.6–32.6), with diversification occurring mainly in the Upper Miocene, although some species groups diversified in the Pliocene and Pleistocene. Biogeographic analysis suggests that the Neotropical *Melanastera* originated from the Pacific region of South and Central America. We detected a shift in diversification rates that likely occurred either at the time of origin of *Melanastera* or during the main colonisation of the Atlantic and Amazon Forests, followed by a subsequent slowdown in speciation rates. State-dependent speciation and extinction models revealed a significant relationship between this diversification shift and the shift of *Melanastera* to the plant families Melastomataceae and Annonaceae, reflecting the impact of host switching on speciation rates in this group. This period also coincides with several independent dispersal events from the Atlantic and Amazon Forests to other parts of the Neotropics. Taken together, the results of the current study suggest that diversification of *Melanastera* was facilitated by major shifts to new host families, which may have promoted the dispersal of *Melanastera* into new adaptive zones with subsequent processes of local speciation.

## 1. Introduction

Deciphering the evolutionary processes that led to the diversification of insects, the most species-rich group of organisms on Earth, is an important step towards understanding the origins of today’s biodiversity. It is estimated that almost half of all known insect species feed on living plants, which makes these insects the most successful herbivore group on the planet (Schoonhoven et al., 2005). As most of them are food specialists (Hardy et al., 2020; Nyman, 2010; Tilmon, 2008), it is particularly interesting to know whether and how interactions with their hosts have influenced the evolutionary pathways and speciation of insects.

According to some authors (Becerra, 2003; Hembry et al., 2014), the diversification of phytophagous insects is promoted by antagonistic coevolution with plants. Coevolution is often used synonymously with cospeciation (de Vienne et al., 2013) when referring to strictly contemporaneous events. While most authors infer coevolution to involve a short-term process of reciprocal adaptations between interacting species, cospeciation typically refers to the matching cladogenetic events over a longer evolutionary time period (Althoff et al., 2014; Cruaud and Rasplus, 2016; de Vienne et al., 2013). Recent cophylogenetic studies question whether coevolution over a short time scale leads to reproductive isolation between populations and thus triggers cospeciation between plants and insects (Althoff et al., 2014; Kahnt et al., 2019). It therefore remains questionable whether coevolution via reciprocal adaptations has contributed substantially to the enormous radiation of insects.

The cospeciation concept, on the other hand, has been criticised because there is little evidence and rare cases of contemporaneous cospeciation patterns (de Vienne et al., 2013; Suchan and Alvarez, 2015). Most studies on herbivorous insects revealed incongruent patterns between the phylogenies of interacting species and some analyses showed that host lineages are generally older than their associated herbivores (Burckhardt and Basset, 2000; Cruaud et al., 2012; Janz, 2011; McKenna et al., 2009; Nyman, 2010; Percy et al., 2004; Tilmon, 2008), leading to patterns of asynchronous divergence between host and insect lineages.

Another model to explain diversification processes in herbivores is the adaptive radiation (‘escape and radiate’ hypothesis) (Allio et al., 2021; Braga et al., 2018; Cruaud et al., 2012; Dasmahapatra et al., 2010; Ehrlich and Raven, 1964; Fordyce, 2010; Janz, 2011; Nylin et al., 2018; Percy, 2003; Thompson, 1994; Wilson et al., 2012). According to this hypothesis, host switches between more distantly related plant lineages open up new adaptive zones for herbivores by providing new resources without competitors and thus triggering rapid species diversification (Janz and Nylin, 2008; Suchan and Alvarez, 2015). In this scenario, adaptations to new hosts are often hypothesised to correlate with the genetic ability to detoxify secondary plant compounds, offering further potential for adaptive radiation in herbivores (Allio et al., 2021; Condamine et al., 2018; Janz, 2011; Nallu et al., 2018). Another interpretation, the ‘oscillation’ hypothesis, suggests slower but constant rates of insect speciation driven by repeated host range expansion (polyphagy) and contraction (monophagy) followed by colonization of new hosts (Hardy, 2017; Janz and Nylin, 2008). Similar to the ‘oscillation’ hypothesis, the ‘musical chairs’ hypothesis suggests that speciation is driven by host switching at a constant rate, but without changes in trophic niche breadth (Hardy and Otto, 2014).

Biogeographical factors are another driving force behind speciation. Phylogenetic studies in the Neotropics have shown that many taxa evolved and diversified during times of major landscape (Neogene: 23–2.6 Ma) or climatic changes (Eocene: 56–33.9 Ma; Pleistocene: < 2.6 Ma) (Chazot et al., 2021, 2019b; Garzón-Orduña et al., 2014; Gentry, 1982; Hoorn et al., 2010; Matos-Maraví, 2016; Rull, 2014; Smith et al., 2014; Matos-Maraví et al., 2021a). The Neotropics is considered the most species-rich biogeographical realm and is often referred to as the cradle or museum of diversity (Antonelli et al., 2018; Fischer, 1960; McKenna and Farrell, 2006; Moreau and Bell, 2013). The ‘cradle’ model postulates that existing species diversity is due to a recent, rapid accumulation of species resulting from high speciation rates, whereas the ‘museum’ model predicts that present species richness results from a steady accumulation of species with constant speciation rates and/or low extinction rates over time (Chazot et al., 2021; Condamine et al., 2012; Lisa De-Silva et al., 2017; Sahoo et al., 2017; Sánchez-Herrera et al., 2020; Toussaint et al., 2019; Winkler et al., 2018). A considerable number of studies on tropical insect–plant systems exploring drivers of species diversity are hampered by small sampling sizes of taxa or limited geographical coverage. Moreover, many of these studies use examples from the Lepidoptera or Hymenoptera groups, but few address groups of plant-sap-sucking Hemiptera.

A potentially useful model taxon for testing these hypotheses are the jumping plant lice or psyllids (Hemiptera: Sternorrhyncha: Psylloidea). Most of the slightly more than 4,000 described species are monophagous or narrowly oligophagous, and related psyllid species tend to develop on related plant taxa (Burckhardt et al., 2021, 2014). The majority of psyllid hosts are eudicots and magnoliids with only a few species living on monocots and less than a handful on conifers (Burckhardt et al., 2014; Hodkinson, 2009; Ouvrard et al., 2015). A number of psyllid species have a narrow distributional range, especially on islands or in mountains (Bastin et al., 2023; Burckhardt, 2024). The true psyllids (Psylloidea s.str.) constitute a relatively young superfamily. Their fossil record is sparse compared to that of other insects (Drohojowska et al., 2020). The oldest known representative, *Eogyropsylla paveloctogenarius* (Aphalaridae), is an Eocene compression fossil from the oil shales of the Kishenehn Formation in northwestern Montana, USA (Ouvrard et al., 2013). With 69 described species (67 extant, 2 fossil), *Melanastera* (Liviidae: Liviinae: Paurocephalini) is the most species-rich genus of Neotropical Psylloidea (Burckhardt et al., 2024c, 2024a; Serbina et al., 2024), followed by the genera *Calinda* (Triozidae), *Calophya* (Calophyidae) and *Mitrapsylla* (Psyllidae) with about 50 species each. The large polyphyletic genus *Trioza* is not considered here. The monophyly of *Melanastera* is supported by both molecular and morphological characters (Burckhardt et al., 2024b). The majority of *Melanastera* species occur in Brazil (60 spp.) (Serbina et al., 2024), with a further nine described species in the New World (7 extant spp., 2 fossil spp.) (Burckhardt et al., 2024b, 2024a) and two in the Old World (Burckhardt et al., 2024c; He et al., 2024). A detailed taxonomic treatment of the Brazilian species was recently published by Serbina et al. (2024). The Neotropical *Melanastera* species are monophagous or narrowly oligophagous on Melastomataceae (Myrtales: one of the basal clades of rosids according to Zuntini et al. 2024; eudicots), Annonaceae (Magnoliales, magnoliids), Asteraceae (Asterales, campanulids, asterids, eudicots), Myristicaceae (Magnoliales, magnoliids) and Cannabaceae (Rosales, fabids, rosids, eudicots), while the two Old World species are associated with Malvaceae (Malvales, malvids, rosids, eudicots), a host family also found in other species of the Paurocephalini (Burckhardt et al., 2024c). Within Melastomataceae, *Miconia* is an extremely diverse and taxonomically difficult genus that occurs throughout the tropical and subtropical Neotropics (Goldenberg et al., 2008; Michelangeli et al., 2022; Ulloa Ulloa et al., 2022). More than one third of all *Melanastera* species (28 of 69 extant spp.) develop on *Miconia* species (Serbina et al., 2024), indicating a major radiation of psyllids on this plant genus.

In this study, we reconstruct a time-calibrated phylogeny of *Melanastera* species using multi-locus DNA sequence data, diversification approaches and fossil records. The resulting phylogenies are used 1) to date the onset of diversification of *Melanastera* in the Neotropics, 2) to assess diversification rates during its evolutionary history, 3) to reconstruct the ancestral host associations and ancestral area of *Melanastera*, 4) to estimate the frequency of host shifts between distantly related plant taxa in the evolution of *Melanastera*, and 5) to analyse the historical biogeography of *Melanastera* species. We test 1) whether *Melanastera* and its associated host plant taxa are of the same age; 2) the significance of cospeciation with hosts, shifts between host families and shifts within host families; 3) the role of vicariance versus dispersal events for explaining present-day distributional patterns, and 4) whether the diversification rates of *Melanastera* accelerated following the colonization of new geographic areas or host families, consistent with the ‘escape and radiate’ and ‘cradle’ hypotheses.

## 2. Material and methods

### 2.1. Psyllid sampling

We analysed 67 extant species of *Melanastera* from Brazil, Belize, Costa Rica, Ecuador and Venezuela (Table S1), six of which are undescribed. Species delimitation was based on morphology, an uncorrected *p*-distance threshold for two barcoding genes, cytochrome oxidase subunit I (*COI*) and cytochrome b (*cytb*), as well as host and distribution data (Serbina et al., 2024). As outgroup taxa, we used *Lanthanaphalara mira* (Aphalaridae), *Euphyllura canariensis* (Liviidae: Euphyllurinae) and several species of Liviinae (*Camarotoscena speciosa*, *Diclidophlebia fremontiae*, *Klyveria crassiflagellata*, *K. flaviae* and *K. setinervis*) which are closely or distantly related to *Melanastera* based on our previous studies (Burckhardt et al., 2024b) and Percy et al. (2018).

Most of the material was collected by D. Burckhardt and D. L. Queiroz from 2011–2024 in 15 states of Brazil (Serbina et al., 2024) and is deposited at the Naturhistorisches Museum, Basel, Switzerland (NHMB) and the Universidade Federal do Paraná, Curitiba, Brazil (UFPR). Additional specimens from Belize, Brazil, Costa Rica, Ecuador and Venezuela were examined from the Natural History Museum, London, UK (BMNH) and the collection of D. M. Percy, Vancouver, Canada. The specimens were swept from the plants with entomological nets on a long stick and collected with an aspirator. Some immatures were collected directly from the plant with an aspirator. The material was immediately preserved in 70% or 96% ethanol for molecular analyses. Voucher specimens were mounted on permanent slides in Canada balsam and are deposited in the entomological collections of the BMNH, NHMB, UFPR and the Moravian Museum, Brno, Czech Republic (MMBC).

### 2.2. Molecular analysis

Most of the sequences of mitochondrial genes *COI* and *cytb* for the current study were taken from Serbina *et al*. (2024). We obtained here new partial sequences of two mitochondrial (*12S rRNA*, *16S rRNA*) and three nuclear (*histone H3*, *wingless* (*wg*), *28S rRNA*) gene regions for 64 *Melanastera* species (Table S1). The molecular sequences for the remaining three *Melanastera* species (*M. lucens*, *M. smithi* and *M.* Venezuela) and most outgroup taxa were taken from Burckhardt et al. (2024b), with the exception of *Lanthanaphalara mira*, whose mitochondrial sequences were obtained from Percy et al. (2018). The primer sequences for the sequenced gene regions, the amplicon sizes and the conditions for PCR used for amplification are as given in Burckhardt et al. (2024b) and here in the Supplementary material (Table S2).

The dry-mounted material from museum collections was stored for up to 28 years prior to DNA extraction (the oldest samples from which we successfully amplified PCR products were collected in 1990). To increase the yield of DNA, we worked with small elution volumes (e.g. 10–20 μL) following Suchan *et al*. (2016) for samples with a low DNA concentration, small specimens and dry-mounted specimens from museum collections. To amplify DNA in low concentrations, we also used nested PCR amplification, which allows amplification of large gene fragments (crucial for *12S rRNA* and *16S rRNA*) and those present in low quantity.

Amplified products were purified using ExoSAP-IT Express (Applied Biosystems) and sequenced in both directions with a BigDye Terminator v3.1 Sequence Kit (Applied Biosystems). The purified PCR products were sequenced on an ABI Prism 3130 Genetic Analyzer (Applied Biosystems). The sequenced fragments were assembled using Sequencher 4.8. The protein-coding gene sequences (*COI*, *cytb*, *H3*, *wg*) were manually aligned using MEGA X (Kumar et al., 2018). The non-protein-coding gene sequences (*12S rRNA*, *16S rRNA*, *28S rRNA*) were aligned with MAFFT v7 (Katoh et al., 2019) using the G-INS-I algorithm (*16S rRNA*, *28S rRNA*) and the Q-INS-I algorithm (*12S rRNA*). All analysed gene fragments were submitted to GenBank (Clark et al., 2016) (Table S1). The program Gblock (Castresana, 2000) with least stringent selection options was used to remove ambiguously aligned blocks in the non-protein coding gene regions.

### 2.3. Phylogenetic reconstruction and estimation of divergence time

All gene sequences were assembled using SEAVIEW v3.2 (Gouy et al., 2010). The codon position for protein-coding gene fragments was set using Mesquite v3.61 (Maddison and Maddison, 2019). The alignment was divided into eleven partitions according to gene regions and codon positions, allowing each partition to have its own evolutionary rate. A phylogenetic analysis was performed using the maximum likelihood (ML) criterion and Bayesian inference (BI).

The ML analysis was performed for standard partitions with IQ-TREE 2 v2.0.6 (Chernomor et al., 2016; Crotty et al., 2019; Trifinopoulos et al., 2016). For molecular dating, we used the least squares dating method (LSD2) implemented in IQ-TREE (To et al., 2016). The 95% confidence intervals were inferred by resampling the branch lengths 1,000 times. The support values for the best-scoring ML tree were determined by performing 1,000 standard bootstrap replicates. The best ML tree was used as a starting topology. Nodes on the final tree with bootstrap support values (BS) > 70% were considered significantly supported, following Hillis and Bull (1993). Divergence times were estimated using the fossil constraints and a lognormal relaxed clock. For the root of the entire tree, we used the age range 47.8–41.3 Ma (Eocene), based on *Eogyropsylla paveloctogenarius* (Aphalaridae: Aphalarinae: Paleopsylloidini†), the oldest known representative of the Psylloidea s.str. (Ouvrard et al., 2013). This fossil was used to calibrate the root of the phylogenetic tree defined by the outgroup *Lanthanaphalara mira*, an extant member of the same subfamily. An additional calibration point was included for dating the stem node of *Melanastera* using the fossil *Diclidophlebia venosa* from Dominican amber (20.4– 13.8 Ma, Lower Miocene) (Burckhardt et al., 2024a), a congener of the extant *D. fremontiae*, and representative of the Paurocephalini (Liviinae: Liviidae) (Fig. 1). We did not use the fossils *Melanastera casca* or *M. vetus* which have the same age (i.e. from Dominican amber: Burckhardt et al., 2024a), to calibrate the crown node of *Melanastera*, as this may lead to an overestimation of the crown age (Chazot et al., 2019a).

**Fig. 1.**
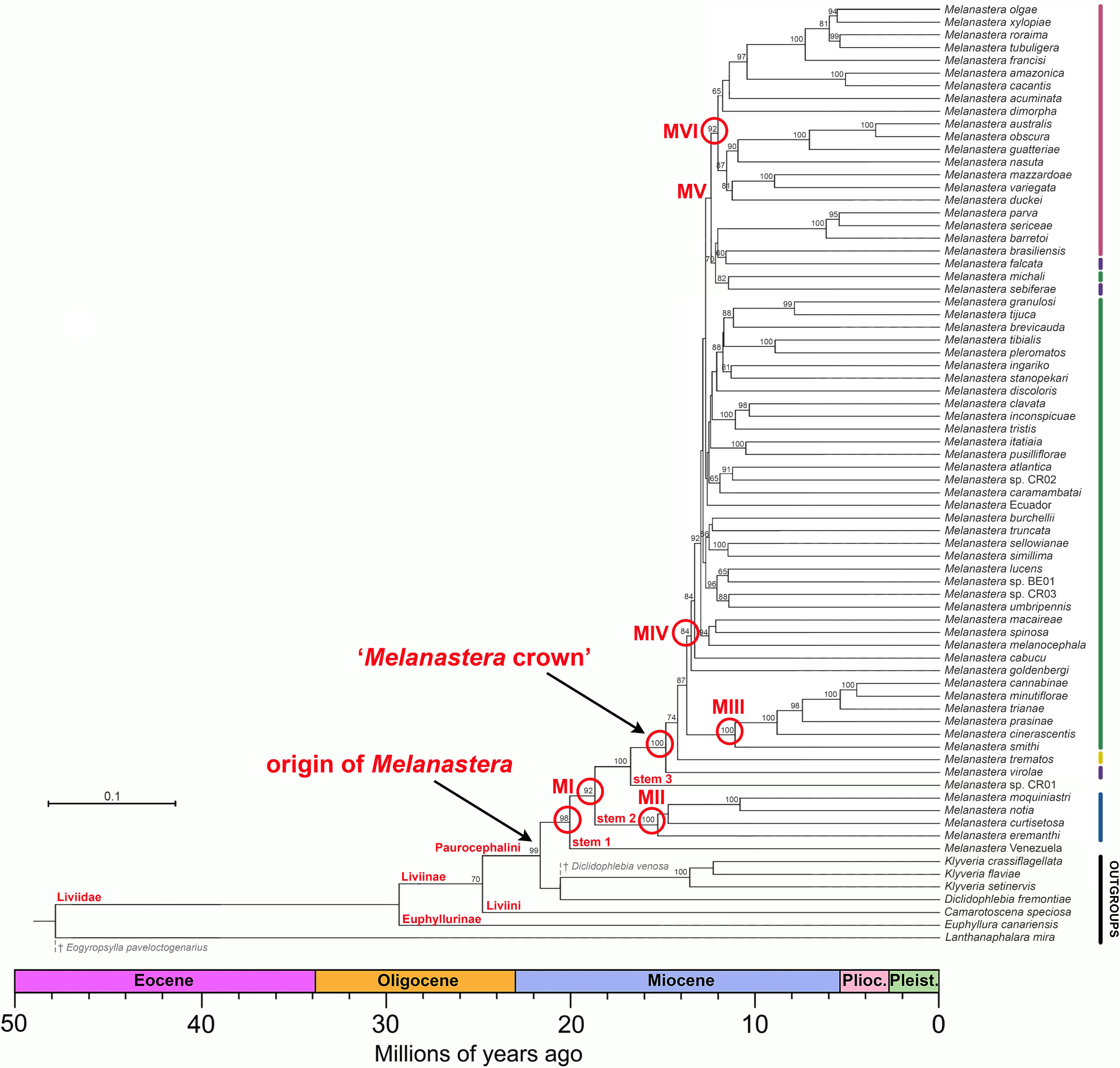
Time-calibrated phylogenetic tree for the genus *Melanastera* with outgroups, based on maximum likelihood analysis of molecular characters. Bootstrap support values (> 70%) are shown next to the crown node. The main clades are marked with red circles; the corresponding age priors can be found in Table S5. Fossil calibration points indicated by grey dashed lines. Coloured bars on the right denote confirmed or potential host family associations.

The BI analysis was performed with BEAST v2.7.5 (Bouckaert et al., 2019; Drummond and Rambaut, 2007). The best substitution model for each partition was estimated using the model-averaging approach implemented in the bModelTest package (Bouckaert and Drummond, 2017). The analysis of 150 million generations, with trees sampled every 1,000 generations, was run on the CIPRES platform (Miller et al., 2010). Random starting trees were assigned to each partition. An uncorrelated lognormal relaxed clock model was assigned separately for each partition. This is a flexible approach for incorporating calibrations that accounts for rate variation between lineages (Drummond et al., 2006). For the estimation of divergence times, we used the package CladeAge (Matschiner et al., 2017). This approach accounts for increased uncertainty in fossil calibrations, which is particularly important for psyllids, as very few fossils are known for this group. It estimates divergence times based on information on the oldest fossil of a clade in combination with estimates of fossil sampling and diversification rates, net diversification (speciation (λ) – extinction (µ)) and turnover (µ/λ) rates, thus overcoming the main problems of node dating (Matschiner, 2019). The speciation and extinction rates were calculated based on the Bayesian analysis of macroevolutionary mixtures (BAMM) as described below. The net diversification (λ–µ) and turnover (µ/λ) rates were set to be within 0.09–0.10 and 0.04– 0.47, respectively. The fossil sampling rates were calculated using the formula r = q[l - F(f)]2/ {[f(0)][l-F(2f)]}, where q is the extinction rate (µ), F(f) is the cumulative distribution function, f(0) is the density function and t is the time interval (Foote, 1997). The fossil sampling rates were set to 0.04–0.16 based on Foote (1997).

Prior distributions for the two calibrated nodes were set to a lognormal prior constrained by the minimum and maximum ages, using the fossils and age ranges as described above.

To select the tree prior (Yule vs. birth–death models), we analysed each type of prior using the marginal likelihood estimates (MLE) of the stepping-stone analysis (Baele et al., 2012; Xie et al., 2011). The marginal likelihood of both tree priors was computed with 1,000 path steps, and the chains running for 1 million generations with log likelihood sampling every 1,000 cycles. This analysis was performed in the older version of BEAST v1.10.4 (Suchard et al., 2018), where this function is available. The Bayes factor (BF) related to the model with the highest-likelihood was calculated as twice its natural logarithm (2lnBF). We considered BF values >10 as significantly favouring one model over another (Kass and Raftery, 1995). The remaining prior parameters for both the Yule and birth–death model trees were left unchanged. The results were then evaluated using Tracer v1.7.1 (Rambaut et al., 2018) using the effective sampling size criterion (ESS > 200); the first 30% of samples were discarded as burn-in. Convergence of relaxed clock parameters and tree height estimates were difficult to achieve and even after 150 million generations some parameters still did not reach the target values. The tree files were combined using LogCombiner v2.4.5 (included in the BEAST package) and the parameter values were annotated to Maximum Clade Credibility (MCC) using TreeAnnotator v2.4.5 (included in the BEAST package). In the BI consensus tree, clades found with posterior probabilities (PP) > 0.90 were regarded as significantly supported, while clades between 0.80 and 0.90 were considered moderately well supported.

Both BI and ML trees were visualized using FigTree v1.4.4 (http://tree.bio.ed.ac.uk/software/figtree/). We also used the programme RogueNaRok (Aberer et al., 2013) to check for the presence of taxa with an uncertain position in the tree that may have caused artificial lowering of branch support. The reconstructed maximum-clade credibility (MCC) tree recovered from the BI analysis was used for the downstream biogeographical and diversification analyses.

### 2.4. Diversification rate analyses

All diversification rate analyses were performed using the time-calibrated MCC tree from the BI analysis. To reconstruct the potential diversification rates in the *Melanastera* clades, we performed Bayesian analysis of macroevolutionary mixtures in BAMM v2.5.0 (Rabosky, 2014), over the entire phylogeny. We ran 10 million generations and sampled every 1,000 generations with the first 20% of samples discarded as burn-in. The R package BAMMtools v2.1.6 (Rabosky et al., 2014) was used to investigate the dynamics of evolutionary rates among *Melanastera* clades that differ in their host associations and to estimate diversification rates across the entire phylogeny of the group over the evolutionary time scale. We used different values (0.1, 1, 2, 5 and 10) of the compound Poisson prior to assess shift probabilities along tree branches and to ensure that the posterior distribution is independent of the prior distribution (Moore et al., 2016). The best-fit shift configuration and the 95% credible set of distinct shifts were calculated using the posterior distribution of the BAMM analysis.

To account for the potential influence of incomplete taxon sampling on the statistical results (Pybus and Harvey, 2000), we incorporated information on unsampled host group diversity and species discovery rates in well-sampled regions, as reported by Serbina et al. (2024). Based on these estimates, we determined that up to half of the species in *Melanastera* could remain unsampled, and therefore we applied a global sampling fraction of 0.5. The AIC, ΔAIC and Akaike weights were applied to interpret how well the different models fit the tree (Burnham and Anderson, 2004). The lowest AIC value indicates the model that best approximates the data. The differences in AIC scores between the models (ΔAIC) were calculated by deduction of AIC of the best constant rate/simplest model from the AIC of the best alternative model. Following Rabosky (2006), the constant rate/simplest model can only be rejected if ΔAIC > 4 for small phylogenies (n = 30) and ΔAIC > 5.5 for large phylogenies (n = 100).

The possible heterogeneity of diversification rates was also investigated by examining the spectral density profile of *Melanastera* constructed from the modified graph Laplacian model (MGL) (Lewitus and Morlon, 2016) of the *Melanastera* tree without outgroups. The spectral density profile and eigenvalues were determined using the ‘*spectR*’ function implemented in the R package RPANDA 2.2 (Morlon et al., 2016). The significance of the modalities, representing the degree of heterogeneity in the diversification rates was assessed using the ‘*BICompare*’ function. The statistical significance of the BIC values for this number of modalities was tested by comparing the corresponding values of the empirical tree and the random bifurcating trees. To investigate the possible time-dependence of diversification rates, we tested the fit of six models (Morlon et al., 2011): two models with constant rates (Yule and birth–death) with four time-dependent models with variation in speciation and extinction rates and in the period of expanding and declining diversity. These models were fitted using the ‘*fit_bd*’ function implemented in the R package RPANDA v2.2. and evaluated using the likelihood-based method. The time dependence was modelled using an exponential function.

To investigate the rate of lineage diversification, we estimated the gamma statistics (γ), which indicates the extent to which the internal nodes of the phylogenetic tree are closer to the root (γ < 0) or to the tips (γ > 0) of the tree than expected for constant rate models (γ = 0). While the negative γ-values support a model of early diversification, suggesting adaptive radiation, the positive γ-values indicate a late burst of speciation or early extinction. We accounted for the effect of incomplete taxon sampling on the diversification rates by using Monte Carlo Constant Rate test (MCCR) and observed distribution of the γ-statistics according to Pybus and Harvey (2000) where a simulated complete phylogeny was also estimated with double the number of extant taxa sampled. To graphically represent patterns of lineage diversification through time, we reconstructed Lineage Through Time plots (LTT), generated under pure–birth (Yule) model using 1,000 trees with 67 *Melanastera* species and outgroup taxa. These trees were then used to construct a mean LTT curve with 95% CI and compared with the empirical LTT curve. In addition, we assessed the fit of four models of clade accumulation to branching times of the phylogeny using the likelihood-based method. Specifically, we compared two constant rate models (Yule and birth–death) with two density-dependent rate models: density-dependent exponential (DDX) and density-dependent logistic (DDL) models (Rabosky and Lovette, 2008). The MCCR test, gamma statistics, LTT plots visualisation and fitting of the diversification model were performed using the R packages laser v2.4.1 (Rabosky, 2007), Ape 5.7-1 (Paradis et al., 2004), treeSim v2.2 (Stadler, 2011), and phytools v1.9-16 (Revell, 2012).

To determine the relationships between host associations and diversification of *Melanastera*, we used the Binary State Speciation and Extinction (BiSSE) (Maddison et al., 2007) and Multiple State Speciation Extinction (MuSSE) (FitzJohn et al., 2009) models, both implemented in the R package diversitree 0.9-10 (FitzJohn, 2012). Three rates were estimated for each host state: speciation (λ), extinction (µ) and transition rates between host states (q). While BiSSE only considers binary character states, MuSSE allows the implementation of multiple character states. In our study, MuSSE was performed to analyse all five host families of *Melanastera*: Melastomataceae, Annonaceae, Myristicaceae, Cannabaceae and Asteraceae. Several variable and constrained MuSSE and BiSSE models were created in different combinations. After identifying and optimising the best-fit models, we performed an MCMC analysis based on 100,000 steps sampled every 1,000 steps. As some studies emphasised that the results of SSE analyses should be interpreted with caution (Beaulieu and O’Meara, 2016; Rabosky and Goldberg, 2015), we complemented our investigation with the Hidden State Speciation and Extinction model (HiSSE) (Beaulieu and O’Meara, 2016). The latter takes into account the effects of the “hidden” states that influence the diversification processes and that are associated with each observed state in the model. To compare different HiSSE models, including the original BiSSE models, we used the R package hisse 2.1.12 (Beaulieu and O’Meara, 2016). Dual transitions between the observed and hidden traits were excluded and all transition rates were set as equal.

### 2.5. Ancestral host reconstruction

Phylogenetic estimation of ancestral host preferences was performed on a fossil-calibrated BI tree of 43 *Melanastera* and four outgroup species (with host data confirmed by the presence of immature stages) using a Bayesian Binary MCMC (BBM) approach (Ronquist and Huelsenbeck, 2003) in RASP v3.2 software (Yu et al., 2015). The fixed model (Jukes Cantor) was used with an equal variation in rates between sites and the number of maximum states for the ancestral nodes was set to two. MCMC chains were run for one million generations, sampled every 100 generations, with 1,000 generations subsequently discarded as burn-in. The information on the host associations of *Melanastera* and outgroup taxa can be found in Table S1. The ancestral host reconstruction aimed to determine the ancestral host family of extant *Melanastera* species and to estimate the number of shifts between host plant families in each branch. Overall, the host associations were categorised into the following nine character states: (A) Melastomataceae, (B) Annonaceae, (C) Myristicaceae, (D) Cannabaceae, (E) Asteraceae, (F) Malvaceae, (G) Oleaceae, (H) Salicaceae, and (I) Solanaceae. The first five families represent the hosts of the analysed *Melanastera* species and the last four those of the outgroups. Each psyllid species is restricted to a single host family.

### 2.6. Ancestral area reconstruction

Geographic distribution data for the species included in our phylogenetic study were mainly obtained from Brown and Hodkinson (1988), Burckhardt et al. (2005) and Serbina et al. (2024). For undescribed species, the data from the locality labels of the examined specimens were used (Table S1). Where available, the data were retrieved as geographical coordinates, otherwise indicative coordinates were assigned to the records based on the available locality information.

We estimated ancestral areas using the likelihood-based dispersal–extinction cladogenesis (DEC) (Ree and Smith, 2008) and the dispersal–vicariance-like (DIVALIKE) (Ronquist and Sanmartín, 2011) models of distribution range evolution, the model that decouples range evolution from cladogenesis, the BayArea-like (BAYAREALIKE) (Landis et al., 2013), and the same models that account for the founder-event speciation parameter (j). These analyses also indicated colonisation rates between bioregions and potential dispersal and allopatric speciation events. To minimise the excessive number of ancestral areas, the maximum number of areas for ancestral nodes was set to three and we excluded species occurring in ≥ four areas: *Melanastera falcata*, *M. melanocephala*, *M. tubuligera* and *M. xylopiae*. For all models, we set the parameters for dispersal (d) and extinction (e) rates to vary freely. The distribution ranges of the species were coded as presence–absence data. We used the R package BioGeoBEARS v0.2.1 (Matzke, 2014) for the calculations of all models. The models were run based on 100 biogeographical stochastic mappings (BSMs) according to Dupin *et al*. (2017) using a calibrated BI tree derived from the molecular dating analysis described above. The AIC and Akaike weights were applied to interpret how well the different models fit the tree.

The definition of areas for the analysis was based on the biogeographical provinces of the Neotropical and Nearctic regions recognised by Escalante *et al*. (2021) and Morrone *et al*. (2022). The following areas were used in the analysis for the ingroup taxa: (1) Paraná (thereafter, Atlantic Forest), (2) Chacoan, (3) Boreal Brazilian (thereafter, Amazonian Forest), (4) Southeastern Amazonian, (5) Pacific, and (6) Mesoamerican. Two additional areas were added for the outgroup taxa, inhabiting Neotropical and Nearctic regions: South American transition zone (7) for *Lanthanaphalara mira*, and Californian (8) for *Diclidophlebia fremontiae*. The Palaearctic region (9) was defined for the other two outgroup taxa, *Euphyllura canariensis* and *Camarotoscena speciosa*. Based on the combined information from Hoorn *et al*. (2010), Iturralde-Vinent (2006), Iturralde-Vinent and MacPhee (1999) and Montes *et al*. (2015), the following series of seven time periods were implemented in the script, taking into account the main geological events of the area’s position in each time:

1. Tree root – 35 Ma. This period is characterised by the presence of a large marine incursion (Pozo Embayment) in western Amazonia, which limited the dispersal between western and eastern Amazonia.
2. 34 to 32 Ma. According to the GAARlandia hypothesis (Iturralde-Vinent, 2006; Iturralde-Vinent and MacPhee, 1999), the Caribbean and South America were connected by the emerged Greater Antilles and the Aves Ridge, suggesting a temporary land connection and thus favouring the dispersal between these areas.
3. 31 to 25 Ma. This period corresponds to the disappearance of the GAARlandia bridge, which reduced the connectivity between the Caribbean and northern South America, while the disappearance of the Pozo Embayment improved the connectivity between the Andes and the Amazonian region.
4. 25 to 15 Ma. This period is characterised by the formation of mountains in the Central and Northern Andes and the subsequent changes in the Amazonian landscape with the spread of wetlands (Pebas System) that separated Northern Andes and the western Neotropics from the rest of South America. These geological events are considered one of the most significant influences on the diversification of Neotropical insect lineages, as they provided opportunities for vicariant and ecological speciation (Chazot et al., 2019b; Elias et al., 2009; Lisa De-Silva et al., 2017; Matos-Maraví et al., 2021a; Sánchez-Herrera et al., 2020; Toussaint et al., 2019).
5. 15 to 10 Ma. The Pebas System was reduced and transformed into a large number of wetlands. This period also coincides with the early stages of the formation of the Isthmus of Panama, which connects North and South America (Bacon et al., 2015; Jaramillo, 2018; McKenna and Farrell, 2006), facilitating the opportunities for adaptive radiation between these areas.
6. 10 to 7 Ma. The uplift of the Northern Andes led to the fragmentation of the Amazon Forest with the formation of the Acre system, which separated the Amazon by a large water system. This contributed to a further reduction in dispersal possibilities between the Andean and Amazonian regions and affected diversification rates in some groups (Sánchez-Herrera et al., 2020).
7. 7 Ma to present. During this period, the Acre system was replaced by the Amazon River and the Isthmus of Panama was completely formed. From 2.6 Ma onwards, the increasing diversification of some Amazonian groups is explained by the Pleistocene climate fluctuations and ecosystem fragmentation affecting the distribution patterns and population structuring of the Amazonian lineages, i.e. the Quaternary glaciation refugia hypothesis (Garzón-Orduña et al., 2014; Smith et al., 2014; Toussaint et al., 2019; Whitmore and Prance, 1988).

For each time period, a matrix of scaling factors was manually created to set the dispersal rates between areas depending on their geographical position, indicating a large or small distance between these areas and/or the appearance or disappearance of geographical barriers due to the geological events mentioned above. The dispersal multipliers were set to 1.0, 0.5 and 0.1 (easy, medium-difficult and difficult dispersal, respectively) to reflect the probability of connectivity between the areas and the subsequent dispersal possibilities in the given time (Table S3). The adjacency matrices were left unconstrained.

## 3. Results

### 3.1. Phylogenetic reconstruction and divergence time estimations

The applied molecular protocols successfully amplified the gene fragments of most species. The most problematic was the amplification of dry-mounted specimens, which failed for the long gene fragments, i.e. *28S rRNA*, *12S rRNA* and *16S rRNA*. The conservative fragment of *28S rRNA* proved to be the most problematic for amplification: it was only successful in 61 of the 74 species analysed. The most successful gene fragment was *cytb*, which could be amplified in 73 species. The final sequence alignment of seven concatenated genes (after ambiguously aligned blocks were removed with Gblock) contained 3222 characters (Table S4).

While the topology of both ML (Fig. 1) and BI (Fig. S1) trees is similar, node support in the ML tree is considerably higher. The molecular dating analyses of the ML and BI trees yielded similar estimates of divergence times, with maximum age differences between the two methods of 9.0–11.5 Ma at deep phylogenetic levels, while the age estimates for the inner clades of the tree showed minor differences (Table S5). These differences in divergence times between two phylogenetic trees can be explained by the different dating approaches in IQ-TREE and BEAST, tip-dating vs. CladeAge dating. While the CladeAge approach in BEAST is similar to the node-dating approach, where the age of internal nodes in the phylogenetic tree is estimated using prior calibrations (Matschiner, 2019; Matschiner et al., 2017), the tip-dating using LSD2 criterion (IQ-TREE) directly uses the ages of the fossil samples as the terminal taxa (To et al., 2016).

The results on the divergence time estimation in the ML tree (Fig. 1) indicate that the Liviidae split into the two subfamilies, Euphyllurinae and Liviinae (100% BS) around 29.5 Ma (CI 25.4–35.2) in the Lower Oligocene. The Liviinae further split into the two tribes Paurocephalini and Liviini (100% BS) in the Upper Oligocene (25.4 Ma; CI 25.4–35.2). The origin (stem node) of *Melanastera* was estimated to be 21.9 Ma (CI 21.9–31.8) at the beginning of the Miocene. *Melanastera* was recovered as monophyletic with strong support (98% BS) and constitutes the sister taxon of the clade containing *Diclidophlebia fremontiae* and *Klyveria* species (99% BS). Diversification of Neotropical *Melanastera* began in the Lower Miocene at 20.2 Ma (CI 20.2–30.6). *Melanastera* Venezuela was found as the sister taxon (stem group 1) to the remaining Neotropical *Melanastera* species (clade MI, 98% BS). The clade MI was divided into clade MII (100% BS), comprising four species associated with plants of the Asteraceae (stem group 2: *M. moquiniastri*, *M. notia*, *M. eremanthi* and *M. curtisetosa*), and a clade containing *M.* sp. CR01 (stem group 3) and the strongly supported clade of most *Melanastera* species, hereafter called the ‘*Melanastera* crown group’ (100% BS). This was followed by the successive splits of *M. virolae* (100% BS) and *M. trematos* (74% BS). The remainder of the ‘*Melanastera* crown group’ consists of clades MIII (100% BS), associated with Melastomataceae, and MIV (84% BS), comprising several poorly to strongly supported clades associated with Annonaceae, Melastomataceae and Myristicaceae. Clade MIII began to diversify later (11.0 Ma; CI 11.0–24.1) than its sister clade MIV (13.3 Ma; CI 13.3–25.7). The short internal and long terminal branches within clade MIV indicate rapid radiation. This clade comprises almost 80% of the known *Melanastera* species (53). The weakly supported clade MV, which is restricted to the order Magnoliales: Annonaceae and Myristicaceae, began to diversify around 12.1 Ma (CI 12.1–24.9). Within MV, clade MVI, which is associated exclusively with Annonaceae, is strongly supported (92% BS). The remaining species within clade MIV, whose internal phylogenetic relationships are poorly resolved, develop on Melastomataceae.

The comparison between the BayesFactor (BF) values calculated using the marginal likelihood estimates (MLE) of the stepping-stone analysis (MLE of Yule and birth–death models are -67471.91 and -67365.27, respectively) supported the Yule model (BF = -213.27). We therefore present the results of the analysis obtained with the Yule tree prior (Fig. S1). The tree topology obtained from the BI analysis is mostly congruent with that from the ML analysis, although with some differences. The results of the divergence time estimation of the BI tree indicate that the Liviidae split into Euphyllurinae and Liviinae (0.78 PP) around 38.5 Ma (95% HPD 28.4–50.8) in the Upper Eocene. The diversification of the Paurocephalini (0.89 PP) began in the Lower Oligocene around 31.5 Ma (95% HPD 22.8–44.5), when *Melanastera* split from the other genera. As in the ML analysis, the monophyly of Neotropical *Melanastera* was strongly supported (1.00 PP). In contrast to the ML tree, *D. fremontiae* is the sister taxon of *Melanastera* (1.00 PP) and *Klyveria* the sister group (0.99 PP) of *D. fremontiae* and the *Melanastera* clade. The origin (stem node) and the diversification (crown node) of *Melanastera* were estimated slightly earlier than in the ML analysis: at the Middle Oligocene (28.3 Ma; 95% HPD 19.6– 38.6) and at the Oligocene–Miocene boundary (23.2 Ma; 95% HPD 16.6–32.6), respectively. In contrast to the ML tree, *M.* Venezuela (stem group 1) in the BI tree together with the clade MII (stem group 2) forms the sister group (0.99 PP) to the rest of *Melanastera* (1.00 PP) and not to all *Melanastera* species. The latter cuts off several monotypic lineages: *M.* sp. CR01 (stem group 3) (1.00 PP), then *M. virolae* (1.00 PP), and finally *M. trematos* (0.81 PP). *Melanastera virolae* is the most basal lineage of a large and strongly supported clade of the ‘*Melanastera* crown group’ (1.00 PP). Although the diversification of the ‘*Melanastera* crown group’ started around 17.2 Ma (95% HPD 12.2–24.3), i.e. Lower Miocene, its species diversified until the Pleistocene. Similar to the results of the ML analysis, clade MIII is strongly supported (1.00 PP) and its crown node was dated younger (11.0 Ma; 95% HPD 6.6–14.5) than its sister group MIV. Although the BI analysis failed to recover most clades within the MIV, the topology and species composition remain similar to the ML tree. Similarly, the analysis of the divergence time estimation showed that the groups within clade MIV probably diversified rapidly one after the other. This is reflected in the short internal branches associated with nodes that show moderate to low support (e.g. clade MV). Clade MVI is strongly supported (0.97 PP); it comprises exclusively species associated with Annonaceae and diversified by 12.9 Ma (95% HPD 8.5–16.9). The remaining *Melanastera* within MIV are associated with Melastomataceae and form a largely unresolved polytomy (MVII).

### 3.2. Diversification rates in Melanastera

Regardless of the established prior values for the rate shifts (Poisson prior) in the BAMM analyses, the best-fitting shift configuration with the highest posterior probability included a single rate shift at the origin of the entire *Melanastera* clade (Fig. 2). This net increase in diversification rate at this point was found with all priors analysed in BAMM. We present here only the results of the analysis with a prior rate shift of 1, while the results of the analyses based on other priors are presented in Table 1. The mean net speciation rate associated with the best-fit shift is more than twice as high (0.16; 95% HPD: 0.12–0.20) compared to the rest of the *Melanastera* clades (mean net speciation rate 0.07; 95% HPD: 0.03–0.12). Extinction rates were relatively low across the phylogeny: the clade with the best-fitting shift had a similar mean extinction rate of 0.03 (95% HPD: 0.002–0.09) compared to the other clades (0.03; 95% HPD: 0.001–0.1). Although the probabilities are lower (30, 16, 11 and 0.05% of samples in the posterior distribution), three alternative shift configurations were also systematically recovered in the analyses. The best supported alternative shift occurred after the origin of *Melanastera* but slightly before its major diversification (‘*Melanastera* crown group’). The second and third alternative diversification rate increases were both less well supported and detected on the stem of the clade comprising *M. trematos* and all species associated with Melastomataceae, Annonaceae and Myristicaceae (except *M. virolae*), as well as on the stem of the clade of *M.* sp. CR01 and the ‘*Melanastera* crown group’ (Fig. S2). Apart from these shifts, we observed a relatively constant diversification rate in the remaining *Melanastera* clades throughout their evolutionary history.

**Fig. 2.**
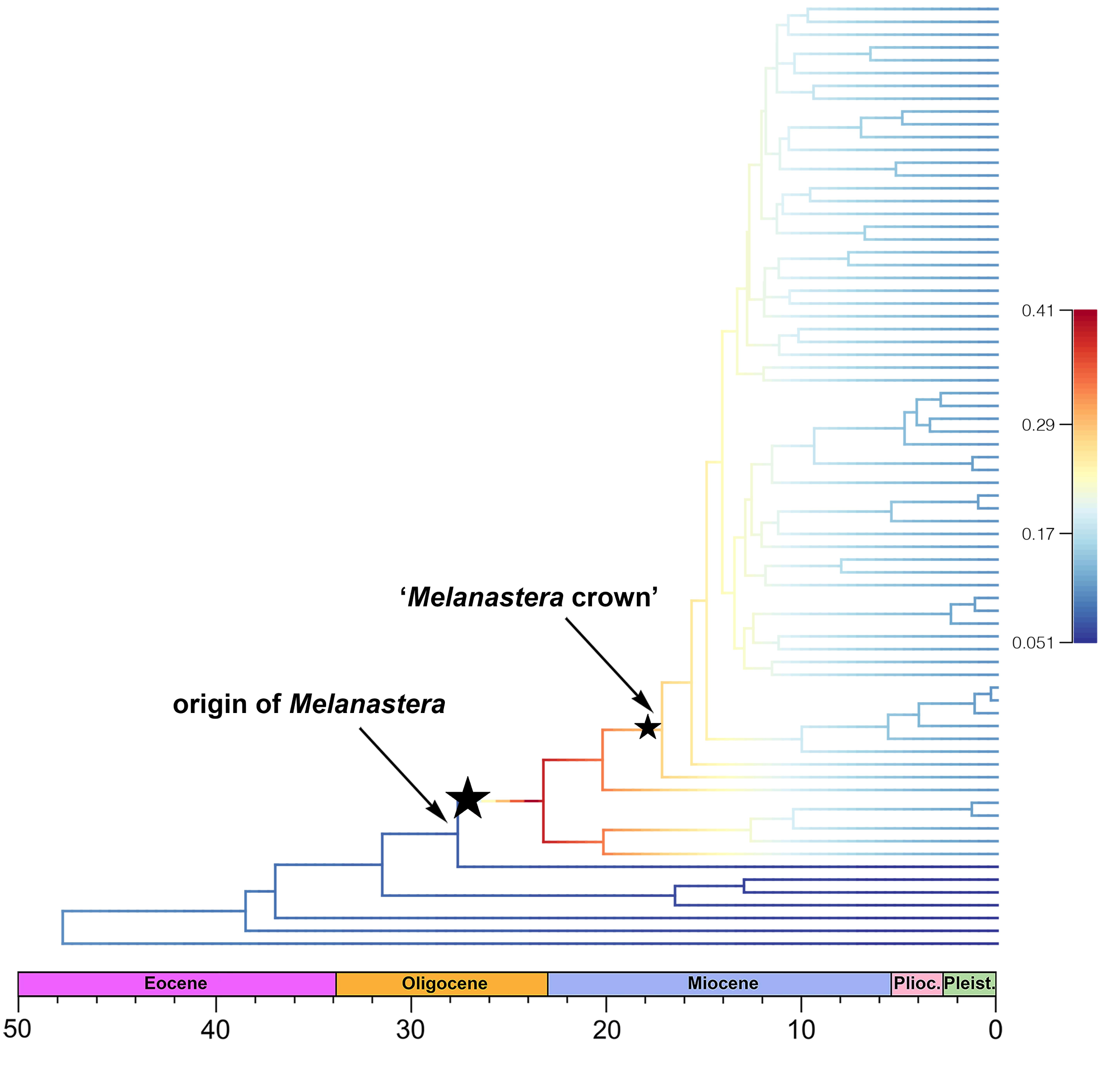
BAMM rate shift configuration with a prior of number of shifts = 1 (see Table 1), based on the phylogenetic tree from Bayesian inference, with branches coloured according to speciation rates. The best-fit (origin of *Melanastera*) and alternative (clade ‘*Melanastera* crown group’) rate shifts are indicated with black stars.

**Table 1.** Results of the BAMM analyses across a gradient of values for the Poisson process governing the number of rate shifts. Prior Shifts – prior on the expected number of shifts; ESS – effective sample size; LnL – log-likelihood; Shift configurations – shift configurations in the credibility set with the probability value (f). The highest probability of shift configuration in the credibility set is in bold.

|  |  |  |  |  |
| --- | --- | --- | --- | --- |
| 0.1 | 10 | 901 | 901 | 1 shift (f = **0.25**), 2 shift (f = 0.23), 3 shift (f = 0.11), 4 shift (f = 0.07), 5 shift (f = 0.03), 6 shift (f = 0.03), 7 shift (f = 0.02), 8 shift (f = 0.01), 9 shift (f = 0.01) |
| 0.2 | 5 | 901 | 901 | 1 shift (f = **0.27**), 2 shift (f = 0.25), 3 shift (f = 0.13), 4 shift (f = 0.09), 5 shift (f = 0.03), 6 shift (f = 0.03), 7 shift (f = 0.02), 8 shift (f = 0.01), 9 shift (f = 0.01) |
| 0.5 | 2 | 901 | 793 | 1 shift (f = **0.35**), 2 shift (f = 0.29), 3 shift (f = 0.16), 4 shift (f = 0.10), 5 shift (f = 0.04), 6 shift (f = 0.03) |
| 1 | 1 | 901 | 793 | 1 shift (f = **0.33**), 2 shift (f = 0.30), 3 shift (f = 0.16), 4 shift (f = 0.11), 5 shift (f = 0.05) |
| 10 | 0.1 | 901 | 680 | 1 shift (f = **0.31**), 2 shift (f = 0.29), 3 shift (f = 0.18), 4 shift (f = 0.16), 5 shift (f = 0.05) |

The BAMM results were also confirmed by constructing the modified graph Laplacian model of the *Melanastera* tree using a model-free analysis of the branching patterns in RPANDA (Fig. S3). The spectral density profile (principal eigenvalue = 8882.98; asymmetry = 0.49; peakedness = 1.73) showed the seven modalities with eigengap between the seventh and eighth of the eigenvalues. The BIC values for the *Melanastera* trees were significantly lower (477,282) than those for randomly bifurcating trees (2,449,593), confirming a significance of the seven modalities of the spectral density profile (BIC ratio = 5.13), which indicated the presence of heterogeneous diversification rates across the *Melanastera* tree.

The temporal variation in diversification rates was confirmed by the ML model-fitting approaches. Among the six Morlon et al. (2011) models tested, the ‘BVAR’ model (exponentially decreasing speciation rate with zero extinction rate) was found to be the best-fitting model with the lowest AIC = 476.21, accounting for almost 70% of the Akaike weight (speciation rate = 0.042 lineages/Ma at present; exponential rate variation = 0.070) (Table 2). Furthermore, the constant rate model was also rejected with certainty due to the ΔAIC, which was > 4. Having found evidence for the time dependence of diversification rates in *Melanastera*, we investigated the possibility that temporal variation may have been influenced by an accumulation of lineage diversity over time. Based on the gamma statistics (γ = –4.67 at *p* = 0.001) and the Monte Carlo Constant rates (MCCR) test (critical γ = –3.18 at *p* = 0.001) which accounts for incomplete taxon sampling, the internal nodes of the tree were significantly close to the root, suggesting a slowdown in the diversification rate in the whole tree. In addition, the lineage through time (LTT) plot deviated from a simulated curve generated assuming a constant diversification rate with incomplete taxon sampling (Fig. 3A). Additional evidence for the slowdown in the diversification rate was obtained by comparing the fit of the diversification models (Yule, birth–death, and density-dependent models). According to the results of this test, the density-dependent logistic model (DDL) was the model with the best support, achieving almost 100% of the Akaike weight. The DDL model (the ‘cradle’ model) was thus favoured over a constant rate diversification scenario (the ‘museum’ model) during the evolution of *Melanastera* (Table 3). Moreover, the constant rate models (Yule and birth–death) were also rejected based on ΔAIC, which was > 5.5. Taken together, the negative value of the gamma statistics (i.e., the position of the internal nodes of the tree near its root) and the best fitting time- and density-dependent diversification models indicated that diversification rates in *Melanastera* varied significantly over time suggesting the pattern of adaptive radiation within the group.

**Fig. 3.**
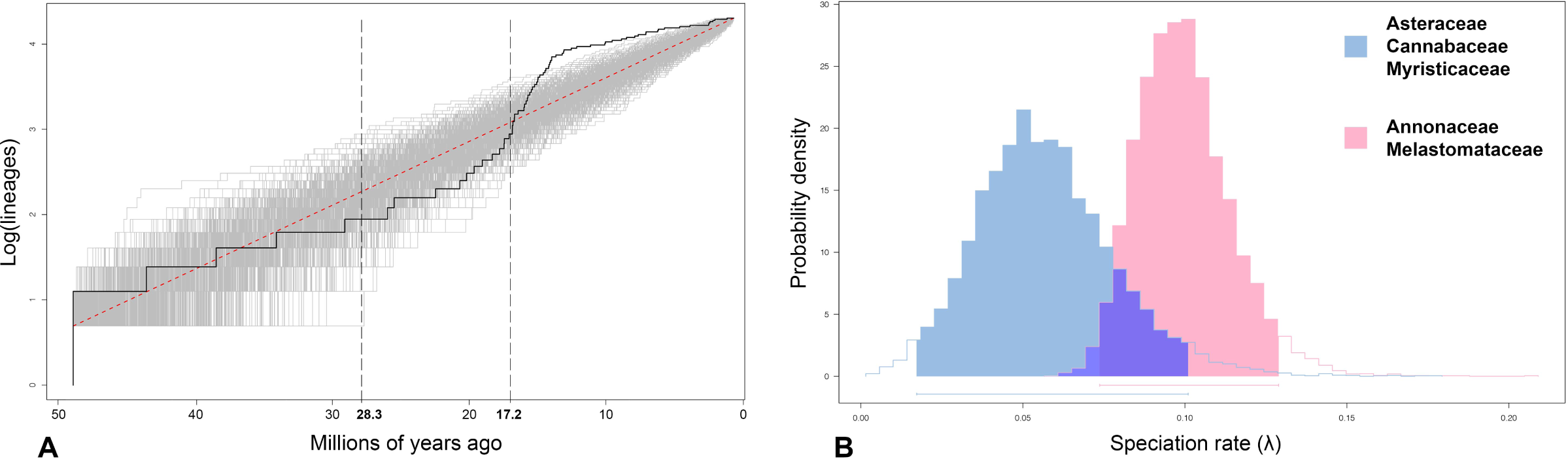
**A.** Lineage through time plot for the genus *Melanastera* with outgroups, based on the Bayesian inference tree. The dashed lines at 28.3 and 17.2 Ma indicate the best and alternative rate shifts determined in the BAMM rate shift configuration, i.e. origin of *Melanastera* (stem node) and ‘*Melanastera* crown group’. **B.** The comparison of speciation rates (λ) between the *Melanastera* lineages associated with the plant families Melastomataceae and Annonaceae vs. Asteraceae, Cannabaceae and Myristicaceae estimated in the best-fit BiSSE model (see Table 4).

**Table 2.** Results of fitting models for the time-dependent lineage diversification. The models are ordered from best (in bold) to worst, by AIC scores and Akaike weights (wtAIC). The ΔAIC scores indicate the difference between the candidate model and the best-fitting model. BVAR = time-dependent speciation; BCST = constant speciation; DCST = constant extinction; DVAR = time-dependent extinction.

| Model | AIC | ΔAIC | wtAIC |
| --- | --- | --- | --- |
| **BVAR** | **476.21** | **0** | **0.69** |
| BVAR - DCST | 478.40 | 2.19 | 0.23 |
| BVAR - DVAR | 480.66 | 4.45 | 0.08 |
| BCST | 487.58 | 11.37 | 0 |
| BCST - DCST | 489.71 | 13.49 | 0 |
| BCST - DVAR | 491.90 | 15.68 | 0 |

**Table 3.** Results of fitting models of lineage diversification of the *Melanastera* tree. The models are ordered from best (in bold) to worst, by AIC scores and Akaike weights (wtAIC). The ΔAIC scores indicate the difference between the candidate model and the best-fitting model. DDL = density-dependent logistic model; DDX = density-dependent exponential model.

| model | AIC | ΔAIC | wtAIC |
| --- | --- | --- | --- |
| **DDL** | **18.18** | **0.00** | **0.999** |
| DDX | 39.96 | 21.78 | 0.001 |
| pure-birth (Yule) | 62.88 | 44.71 | 0.001 |
| birth-death | 64.88 | 46.71 | 0.001 |

**Table 4.**
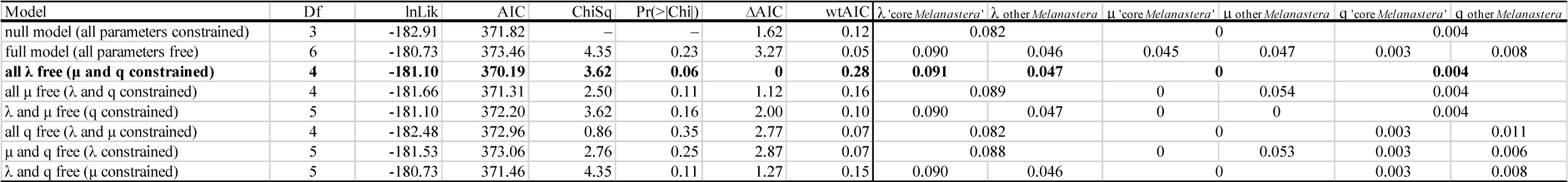
Results of the analysis of variance (ANOVA) of the different models used in the BiSSE analysis to compare diversification rates between *Melanastera* associated with Melastomataceae and Annonaceae (‘*Melanastera* crown group’) vs. *Melanastera* associated with other plant families. Parameter estimates are labelled as follows: λ = speciation rate; μ = extinction rate; q = transition rate between different character states. The abbreviations are as follows: Df = degrees of freedom; lnLik = log-likelihood; AIC = Akaike Information Criterion. The best-fitting model is printed in bold.

The influence of host association on the diversification of *Melanastera* was analysed using the BiSSE model, which only allows binary characters. In agreement with the results of BAMM and RPANDA, we found a difference in speciation rates between *Melanastera* mainly associated with Melastomataceae and Annonaceae and the rest of *Melanastera* associated with other families (i.e. Myristicaceae, Cannabaceae and Asteraceae) (Fig. 3B). The highest speciation rates were estimated for Melastomataceae- and Annonaceae-associated species (λ1 > λ2), whereas *Melanastera* associated with Myristicaceae, Cannabaceae and Asteraceae showed slower speciation rates, with both extinction (μ1/μ2) and transition (q1/q2) rates remaining similar (Table 4). Moreover, transition rates for host switches were often estimated close to zero, indicating a strong host conservatism in *Melanastera*: once a psyllid species switched to a new plant family (Melastomataceae and Annonaceae), it rarely switched back (e.g. to Myristicaceae) or colonised a different family. Although the BiSSE analysis supported the model with varying speciation rates, other models could not be completely discarded as the ΔAIC threshold between all was below 4. Similar to the BiSSE analysis, the HiSSE results were inconclusive, so the presence of “hidden” states that may have influenced the diversification processes of *Melanastera* cannot be excluded (Table 5). Interestingly, when the MuSSE models were applied, the strongest support was suggested for the null model, meaning that all parameters had equal rates (Table 6). This model fitted the data significantly better than most of the other MuSSE models (ΔAIC > 4).

**Table 5.** Results of the analysis of variance (ANOVA) of the different models used in the HiSSE analysis to compare diversification rates between *Melanastera* associated with Melastomataceae and Annonaceae (‘*Melanastera* crown group’) vs. *Melanastera* associated with other plant families. Abbreviations are as follows: lnLik = log-likelihood; AIC = Akaike Information Criterion, CID-2 and CID-4 correspond to the character-independent diversification models, allowing two and four hidden categories, respectively. The best-fitting models are printed in bold.

| Model | lnLik | AIC | ΔAIC | wtAIC |
| --- | --- | --- | --- | --- |
| HiSSE null (trait-independent model, | -186.93 | 379.86 | 2.77 | 0.09 |
| **BiSSE null (trait-independent model,** | **-182.54** | **377.09** | **0** | **0.36** |
| HiSSE (trait-dependent model, with | -186.97 | 383.95 | 6.86 | 0.01 |
| **BiSSE (trait-dependent model, no** | **-182.54** | **377.09** | **0** | **0.36** |
| CID-2 | -186.93 | 378.65 | 1.57 | 0.16 |
| CID-4 | -182.87 | 383.74 | 6.65 | 0.01 |

**Table 6.** Results of the analysis of variance (ANOVA) of the different models used in the MuSSE analysis to compare diversification rates between *Melanastera* associated with Melastomataceae, Annonaceae, Myristicaceae, Cannabaceae and Asteraceae. Parameter estimates are denoted as follows: λ = speciation rate; μ = extinction rate; q = transition rate between different character states. Abbreviations are as follows: Df = degrees of freedom; lnLik = log-likelihood; AIC = Akaike Information Criterion. The best-fitting model is printed in bold.

| Model | Df | lnLik | AIC | ChiSq | Pr(>Chi) | ΔAIC | wtAIC |
| --- | --- | --- | --- | --- | --- | --- | --- |
| **null model (all parameters constrained)** | **3** | **-200.62** | **407.24** | — | — | **0** | **0.84** |
| full model (all parameters free) | 30 | -195.26 | 450.51 | 10.73 | 0.99 | 43.27 | 0 |
| all $\lambda$ free ($\mu$ and q constrained) | 7 | -198.53 | 411.06 | 4.19 | 0.38 | 3.81 | 0.12 |
| all $\mu$ free ($\lambda$ and q constrained) | 7 | -199.84 | 413.68 | 1.56 | 0.82 | 6.44 | 0.03 |
| $\lambda$ and $\mu$ free (q constrained) | 11 | -198.50 | 419.00 | 4.25 | 0.83 | 11.75 | 0 |
| all q free ($\lambda$ and $\mu$ constrained) | 22 | -195.04 | 434.09 | 11.16 | 0.92 | 26.84 | 0 |
| $\mu$ and q free ($\lambda$ constrained) | 26 | -193.98 | 439.95 | 13.29 | 0.94 | 32.71 | 0 |
| $\lambda$ and q free ($\mu$ constrained) | 26 | -192.38 | 436.76 | 16.48 | 0.83 | 29.52 | 0 |

### 3.3. Ancestral host plant associations and host shifts in Melanastera

The reconstruction of ancestral host associations was carried out for 47 *Melanastera* species with known hosts. These species are associated with five plant families (Table S1). The analysis showed that closely related *Melanastera* species tend to develop on hosts of the same family (Fig. 4). However, none of the species groups developing on Annonaceae, on Melastomataceae or on Myristicaceae constitute a monophylum. The only monophyletic group of *Melanastera* species defined by the host plant family is represented by the Asteraceae-feeders (MII, stem group 2), and the group associated with the plant family Cannabaceae is monotypic (i.e. it comprises only one species, *M. trematos*).

**Fig. 4.**
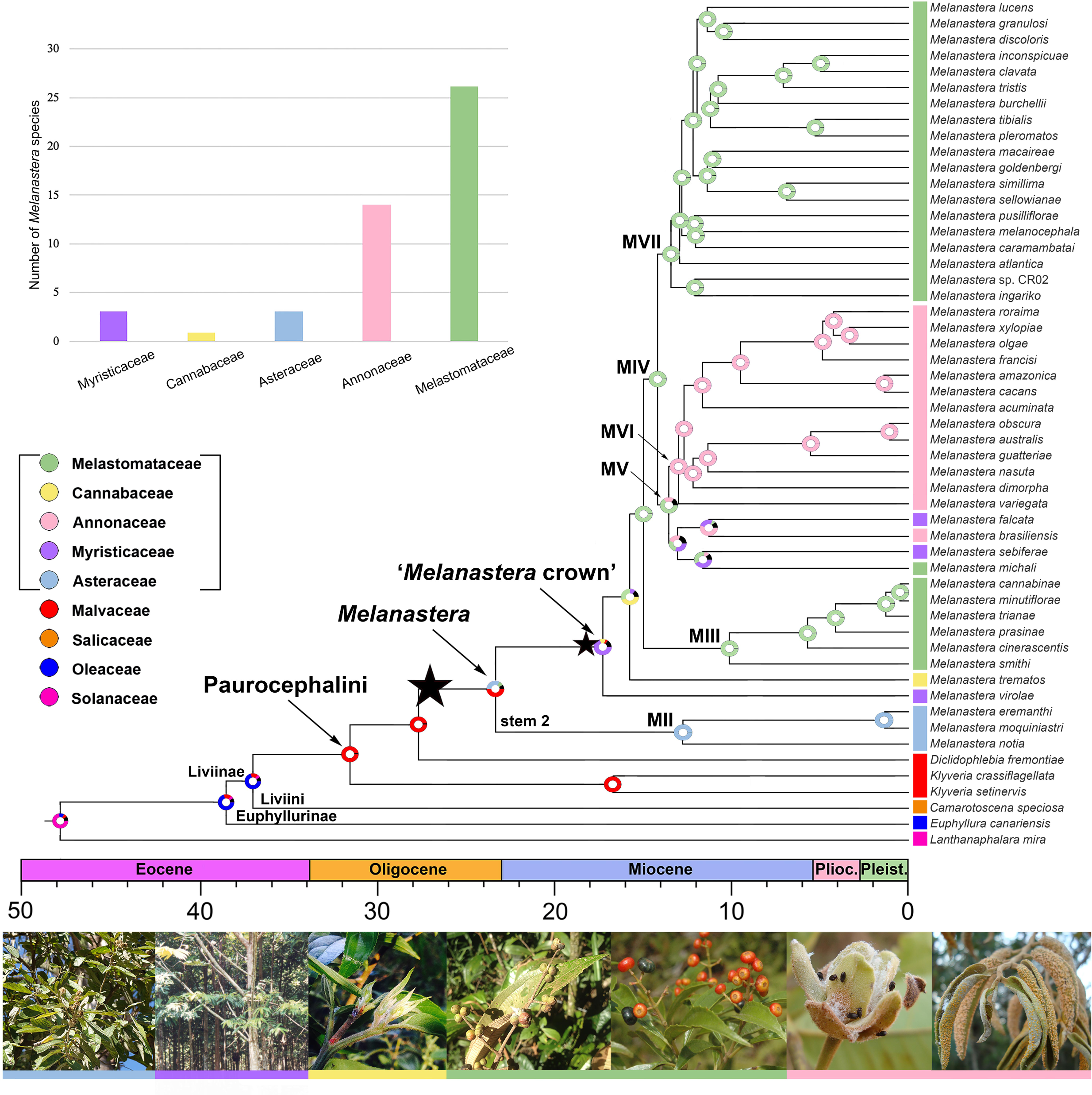
Reconstruction of the ancestral host plant family for the genus *Melanastera* with outgroups, based on the time-calibrated Bayesian inference tree. The best (origin of *Melanastera*) and alternative (clade ‘*Melanastera* crown group’) rate shifts identified in the BAMM analysis are indicated with black stars. Coloured bars on the right denote host family associations. Photographs of representatives of the host plant families can be found at the bottom of the figure, from left to right: *Moquiniastrum polymorphum* (Asteraceae, credit: IM), *Virola sebifera* (Myristicaceae, credit: Jorge Yared), *Trema micrantha* (Cannabaceae, credit: LŠS), *Miconia cinerascens* and *M. sellowiana* (Melastomataceae, credit: LŠS and RG), *Guatteria punctata* and *Xylopia aromatica* (Annonaceae, credit: DQ).

According to the results of the analysis based on the BBM approach, Malvaceae were identified as the most likely ancestral host taxon of the Paurocephalini (0.97 PP). Malvaceae (0.50 PP) or Asteraceae (0.32 PP) appear to be the likely ancestral hosts of *Melanastera*, from which several later host switches to Myristicaceae, Cannabaceae and Melastomataceae may have occurred. The most likely ancestral host of the ‘*Melanastera* crown group’, which unites all *Melanastera* in this analysis except the Asteraceae-associated clade (MII), appears to be Myristicaceae (0.64 PP). Our results suggest at least eight host family changes during the evolutionary history of *Melanastera*. The Melastomataceae, which harbour the most *Melanastera* species (26), were colonised at least once, while the Annonaceae (14 *Melanastera* species) were colonised at least twice independently. A single shift each is suggested for Cannabaceae and Asteraceae, taxa associated with only one and three species, respectively, in the analysis. The Myristicaceae were colonised three times independently, more frequently than the other plant families within *Melanastera*, although only three *Melanastera* species are associated with this family.

### 3.4. Reconstruction of the ancestral area and potential dispersal routes for Melanastera

The results of the historical biogeographic analysis showed that the DIVALIKE+J model had the lowest AIC value (328.5) with 66% of the Akaike weights and the lowest likelihood LnL = -161.6, and was therefore selected as the preferred biogeographic model (Fig. 5). This emphasised the importance of founder event speciation for the evolution of *Melanastera* in addition to dispersal and vicariant events.

**Fig. 5.**
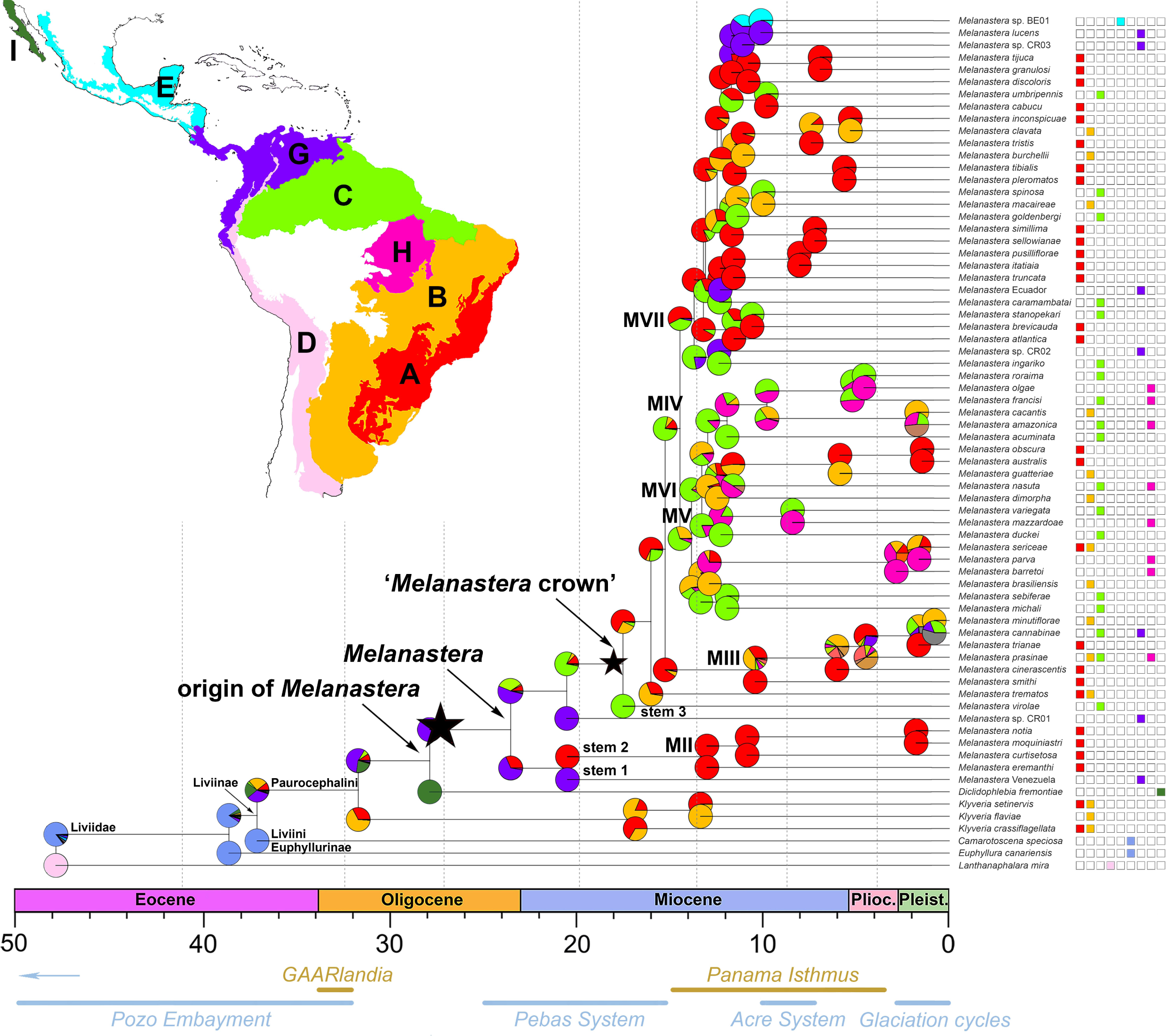
Reconstruction of the ancestral area for the genus *Melanastera* with outgroups, based on the time-calibrated Bayesian inference tree, under the DIVALIKE+J model. The geographical distribution of each species is shown on the right side of the figure. The biogeographical provinces of the Neotropical region are shown on the map (modified from Morrone et al., 2022): A – Paraná (Atlantic Forest), B – Chacoan, C – Boreal Brazilian (Amazon Forest), D – South American transition zone, E – Mesoamerican, G – Pacific, H – Southeastern Amazonian, I – Californian. The best (origin of *Melanastera*) and alternative (clade ‘*Melanastera* crown group’) rate shifts identified in the BAMM analysis are indicated with black stars.

According to the DIVALIKE+J model, the Pacific region (G) had the highest probability of being the area of origin of *Melanastera*. However, in contrast to other areas, the Pacific region comprises a relatively small number of *Melanastera* species (7) which were analysed here. We hypothesise that the first colonisation events from the Pacific region into the Atlantic (A) and Amazon (C) Forests may have started at the beginning of the Miocene, followed by multiple dispersal events into other parts of the Neotropics and a repeated recolonisation of the Atlantic Forest, the Amazon Forest and the Pacific region that continued into the Pleistocene.

The Atlantic Forest was the most frequently colonised independently by *Melanastera* in the course of its evolutionary history. The first dispersal event from the Pacific region into the Atlantic Forest occurred at the beginning of the Miocene, followed by the diversification of clade MII, which represents a small radiation within the Atlantic Forest area. This period also coincides with the diversification of the two species within the Pacific region, namely *M.* sp. CR01 from Costa Rica and *M.* sp. from Venezuela. Interestingly, the two species inhabit the Caribbean and northern South America, which were separated from the rest of South America by the Pebas System (Hoorn et al., 2010), suggesting a vicariant origin of both *M.* sp. CR01 and *M.* Venezuela, independently of each other. The beginning of the Miocene also corresponds to the time of the first dispersal into the Amazon Forest followed by the speciation of *M. virolae*. The next colonisation of the Atlantic Forest did not begin until the Middle Miocene, leading to a rapid radiation of the ‘*Melanastera* crown group’, which was identified as the alternative rate shift point in the diversification analyses in our study. Clade MIII probably originated in the Atlantic Forest and underwent several colonisation events into the Chacoan and the Amazon regions, as well as the reverse dispersal back into the Pacific region.

The next great colonisation was reconstructed from the Atlantic Forest to the Amazon. As a result of these successive colonisation events, we could identify a series of long and short distance dispersal events that allowed the colonisation of new territories, the Chacoan (B) and Southeastern Amazon (H) regions, and a further recolonisation of the Atlantic, Amazonian and Pacific regions. Among them, allopatric speciation can only be assumed for the *M. mazzardoae* – *M. variegata* clade distributed in the Southeastern Amazon and the northern parts of the Amazon Forest, respectively, which were separated by the Acre System at that time (∼8 Ma) (Hoorn et al., 2010). The diversification of several terminal clades, i.e. *M. sericeae* – *parva* – *barretoi*, *M. australis* – *obscura* – *guatteriae*, *M. cacantis* – *amazonica*, *M. roraima* – *olgae* – *francisi*, occurred within different areas (A, B, C and/or H) with no evidence of vicariant speciation.

The poor clade support for the other groups within the MIV makes it impossible to surmise in detail further dispersal routes for *Melanastera*. Nevertheless, we assume a dispersal back to the Atlantic Forest, the Chacoan, the northern Amazon and the Pacific regions, as well as a first colonisation of the Mesoamerican region. The latter probably occurred from the Pacific region and has been estimated at 9.7 Ma, the time of the formation of the Isthmus of Panama, which connects North and South America (Jaramillo, 2018). Interestingly, almost half of the representatives of clade MVII (15 of 29 spp.), which diversified from the end of the Upper Miocene to the Pliocene originated exclusively from the Atlantic Forest.

## 4. Discussion

The adaptive radiation scenario suggests that lineage diversification rates accelerate as a consequence of ecological opportunities, such as the availability of new ecological niches and the lack of natural enemies (Janz and Nylin, 2008; Suchan and Alvarez, 2015). Many studies on herbivore taxa from the Neotropics have shown that it is the radiations following major host switches and dispersal events that particularly drive diversification in these groups (Chazot et al., 2019b; Lisa De-Silva et al., 2017; Sahoo et al., 2017; Sánchez-Herrera et al., 2020; Toussaint et al., 2019). After niches are filled and new opportunities diminish, speciation rates decrease, slowing the pace of lineage diversification (cradle model) (Rabosky, 2014). On the other hand, diversification patterns in some insect groups show constant speciation and extinction rates, arguing in favour of the Neotropics as a museum of diversity (Condamine et al., 2012; Winkler et al., 2018), according to which today’s Neotropical diversity has been shaped by a steady accumulation of species over time. In the present study, we performed several phylogenetic and diversification analyses to understand the evolutionary processes that have shaped the diversification of the psyllid genus *Melanastera* in the Neotropics.

### 4.1. *Systematics of* Melanastera

The taxonomy of Brazilian *Melanastera* species was recently revised based on morphological and molecular characters (Serbina et al., 2024). In the present phylogenetic analyses, 61 (out of 69) described and six undescribed Neotropical species were included. The maximum likelihood analysis (ML) and Bayesian inference (BI) trees produced similar topologies, with generally high nodal support, though some nodes in the central part of the phylogenetic trees were left unresolved. In both analyses, Neotropical *Melanastera* was strongly supported as monophyletic, justifying the genus concept by Burckhardt et al. (2024c). However, the study by Burckhardt et al. (2024c) included a much smaller subset of *Melanastera* species (5) and found only weak to moderate support for the monophyly of *Melanastera* when using combined molecular and morphological datasets. In the ML tree, *Melanastera* Venezuela was strongly supported as a sister taxon to all other Neotropical *Melanastera* species, in contrast to the weaker or moderate support found in previous analyses by Burckhardt et al. (2024b). In contrast, the BI analysis placed *M.* Venezuela as a sister taxon to the MII clade, which includes species associated with the host family Asteraceae. In general, most clades in *Melanastera* form short internal branches that evolved within a relatively short period of time, while the terminal branches remain relatively long. As suggested in other phylogenetic studies, the presence of short branches in the tree may be the result of rapid speciation events in which new species form in a short period of time without fully establishing distinctive species boundaries (Whitfield and Lockhart, 2007). Such rapid speciation can therefore lead to the emergence of numerous species with little morphological and molecular divergence (Nyman et al., 2010; Tilmon, 2008), which also appears to be the case in *Melanastera* (Serbina et al., 2024).

Serbina et al. (2024) defined nine species groups for *Melanastera* based on morphological characters, primarily for practical reasons of species description and identification. They acknowledged that most of these groups are likely artificial. This study confirms their assertion through phylogenetic analyses, with the exception of the *M. curtisetosa* group of Serbina et al. (2024), of which the extant Neotropical members in the current study form a monophylum (clade II). There is also solid support for the monophyly of the *M. smithi* group by Serbina et al. (2024) when *M. sellowianae* and *M. simillima* are excluded (clade MIII in the current study).

### 4.2. Diversification times and host evolution in Melanastera

Hosts of *Melanastera* are reported from five unrelated plant families. The phylogenetic patterns do not indicate a single colonisation of the individual plant families, with the exception of the Asteraceae and Cannabaceae. The Melastomataceae host about 55% of the *Melanastera* species analysed, most of which are associated with the tribe Miconieae (especially the genus *Miconia*) and several species with the tribes Melastomateae (host genus *Pleroma*) and Marcetieae (*Macairea*). The Melastomataceae are an extremely species-rich group with around 5,900 species worldwide, of which around 3,740 occur in the Neotropics (Ulloa Ulloa et al., 2022). The Melastomataceae originated in the Upper Cretaceous, their Neotropical tribes separated in the Eocene and a major radiation of Miconieae in the Neotropics occurred in the Miocene (Maurin et al., 2021; Penneys et al., 2022; Reginato et al., 2022). The origin of Annonaceae, another well-represented host family within *Melanastera*, is also dated to the Cretaceous (Massoni, 2015; Zuntini et al., 2024). This plant family accounts for almost 30% of the host plants of the *Melanastera* species analysed, which are associated with the plant genera *Annona*, *Guatteria* and *Xylopia*. These three genera are not closely related and their common ancestor has been dated to the Palaeocene (ca. 57 Ma) (Couvreur et al., 2011). Our results suggest that *Melanastera* originated at the beginning of the Miocene around 21.9 Ma (ML, CI 21.9–31.8) (Fig. 1) or at the Middle Oligocene 28.3 Ma (BI; 95% HPD 19.6–38.6) (Fig. S1). However, the diversification of most species took place in the Upper Miocene and continued until the Upper Pliocene (ML) or the Upper Pleistocene (BI) (Figs. 1, S1 and 4). The onset of *Melanastera* diversification thus lags well behind the origins of both Melastomataceae and Annonaceae (Massoni, 2015; Zuntini et al., 2024). At the same time, it aligns with the Neotropical radiation of the tribe Miconieae (Michelangeli et al., 2022). This concordance is further supported by the pattern in which most host plants of *Melanastera* associated with Miconieae are restricted to a large clade within the Miconieae phylogeny (Michelangeli et al., 2022). Despite this correlation, the topologies of phylogenetic trees of *Melanastera* and Melastomataceae lack congruence. There is no clear evidence of cospeciation, as the host genera are distributed across different unrelated tribes of Melastomataceae (Fig. S4) and different clades even at the fine level of the phylogeny of Miconieae (Renato Goldenberg, pers. obs.).

Our dataset is biased towards lowland South American forests, with limited representation of *Melanastera* species from the Caribbean and Andes, which are the regions with high *Melastomataceae* diversity (Michelangeli et al., 2022). Another limitation is the low support of some nodes in our *Melanastera* phylogenies. Nevertheless, our results provide a detailed scenario for the diversification of *Melanastera.* We suggest that the extensive radiation of Melastomataceae, particularly Miconieae, with many closely related species, likely facilitated the diversification of *Melanastera*. At the same time, the absence of barriers to host switching in psyllids may have contributed to diffuse, rather than clustered, patterns of host selection from an evolutionary perspective. Similar results emphasising the crucial role of host diversity in diversification have been observed in other groups of herbivorous insects (Forbes et al., 2017; Ward et al., 2022).

The ancestral host family for all *Melanastera* species remains ambiguous. Nevertheless, BBM ancestral reconstructions have shown that Malvaceae has the highest probability of being the ancestral host family of *Melanastera* compared to other families (Fig. 4). Based on the timing of the origin and divergence of Malvaceae in the Upper Cretaceous (∼91 Ma; 95% HPD 69.6–112.3) (Zuntini et al., 2024), it preceded the origin of *Melanastera*. Considering that many other Paurocephalini species, as well as the Old World *Melanastera* species (i.e. *M. pilosa* and *M. sinica*) not included in the current analyses, are indeed associated with Malvaceae (Burckhardt et al., 2024b; He et al., 2024), the ancestral association of *Melanastera* with Malvaceae is even more likely, but future studies including a broader sampling of Paurocephalini representatives are needed to confirm this scenario.

The BAMM converged towards the identify at least one macroevolutionary regime corresponding to an early divergence of *Melanastera*, after which there was a subsequent decline in diversification rates (Figs. 2 and S2). The pattern of heterogeneity in diversification rates was also confirmed by the time- and density-dependent diversification models (Figs. 3 and S3; Tables 2 and 3). During the Miocene, *Melanastera* probably underwent a rapid diversification, a burst triggered by shifts to the new host families Melastomataceae and Annonaceae, leading to an increase in speciation rates, as suggested by our BAMM and BiSSE results (Figs. 2 and 3). Interestingly, not all host shifts were successful from an evolutionary perspective in terms of the extant species diversity of *Melanastera*. Thus, given the high number of *Melanastera* species (40 in the current study) associated with Melastomataceae and Annonaceae, the lineages associated with Asteraceae, Cannabaceae and Myristicaceae are species-poor, each accounting for < 10% of the total diversity (Fig. 4; Table S1). The highly unbalanced extant species richness of Melastomataceae and Annonaceae feeders could be explained by different speciation rates between species associated with these two plant families and those developing on Asteraceae, Cannabaceae and Myristicaceae. While the latter two plant families are relatively species poor and thus unlikely to support significant psyllid radiations, Asteraceae is the most species rich plant family worldwide. Several psyllid groups, such as the genera *Craspedolepta* and *Calinda* show significant radiations on Asteraceae (Loginova, 1963; Olivares and Burckhardt, 1997; Ouvrard et al., 2015). Despite being associated with a relatively large number of plant families, *Melanastera* is characterised by relatively infrequent shifts to new host families, while frequently transitioning between hosts within the same family. This is consistent with the findings of Ouvrard et al. (2015) on the evolution of host associations of psyllids and similar studies on other herbivorous groups, which have shown that most herbivores exhibit strong phylogenetic conservatism in their host associations. They tend to favour shifts to chemically similar and therefore, phylogenetically related and therefore chemically similar plant taxa, while shifts to more distantly related taxa do occur but are relatively rare (Nyman et al., 2010; Suchan and Alvarez, 2015).

In our study, using various time- and density-dependent diversification models, we indeed found a pattern of speciation decreasing with standing diversity of *Melanastera* that supports Ehrlich and Raven’s ‘escape and radiate’ hypothesis, according to which closely related herbivores have similar host preferences (Janz, 2011; Suchan and Alvarez, 2015). This may explain the high species richness of *Melanastera* on plant families such as Melastomataceae and Annonaceae, where new ecological opportunities (i.e. plant species richness) have likely promoted increased speciation rates in *Melanastera* lineages, contributing to the evolutionary success of the group. In contrast to the ‘escape and radiate’ hypothesis, the oscillation and musical chair hypotheses, which also link herbivore diversity to host diversity, suggest more uniform diversification rates (Hardy and Otto, 2014; Janz and Nylin, 2008), which is not consistent with our results. Similar processes contributing to species diversification have been described in other studies on different insect groups associated with species-rich plant groups from the Neotropics (Allio et al., 2021; Jahner et al., 2017; Sahoo et al., 2017; Toussaint et al., 2019; Winter et al., 2017).

### 4.3. Historical biogeography of Melanastera

Most species of *Melanastera* occur in the tropics and subtropics of the New World, extending from Mexico in the north to southern Brazil in the south (Burckhardt et al., 2024b, 2024a; Serbina et al., 2024). Only two species are known from the Old World (Burckhardt et al., 2024b; He et al., 2024).

In the New World, *Melanastera* most likely originated in the Pacific region (Fig. 5) at the boundary between the Oligocene and Miocene. This region includes part of the Caribbean and northern South America, which were not connected to North America by the Isthmus of Panama at the time of the origin of *Melanastera* (Bacon et al., 2015; Jaramillo, 2018), while a temporary connection through the Greater Antilles and the Aves Ridge has already disappeared (Hoorn et al., 2010).

Surprisingly, in this study we found only a few instances of vicariant speciation in *Melanastera*. One of these events coincides with the timing of the earliest colonisation of *Melanastera* (*M.* Venezuela and *M.* sp. CR01, independently) from the Pacific region into the Amazon Forest, which probably occurred in the Lower Miocene (∼20.2 Ma) before the formation of the Pebas System (25–15 Ma). During this time, the Pebas System separated the western part of the Pacific region from its eastern part and from the rest of Amazonia (Hoorn et al., 2010). As shown in other studies, this geological event hindered the dispersal of species and allowed allopatric speciation between these areas and was therefore very important for the speciation of a number of clades of Neotropical butterflies (Chazot et al., 2019b; Condamine et al., 2012; Lisa De-Silva et al., 2017; Matos-Maraví et al., 2021b; Toussaint et al., 2019). Another example of vicariant speciation (*M. mazzardoae* – *M. variegata* occurred during the formation of the Acre System in western Amazonia in the Upper Miocene (10–7 Ma), which separated most of the Amazon Forest from the more southern parts of South America (Hoorn et al., 2010; Sánchez-Herrera et al., 2020). The few cases of potential vicariant speciation in *Melanastera* may be due to incomplete taxon sampling, as many *Melanastera* species probably remain undescribed (Serbina et al., 2024) or were not included in the current analyses. In addition, *Melanastera* was only collected in 15 of 26 Brazilian states, with varying levels of sampling effort in each state (Serbina et al., 2024). These factors may also hinder the accurate interpretation of vicariant events in the evolutionary history of Neotropical *Melanastera*.

Nevertheless, our results suggest that the diversification of *Melanastera* into different lineages was mainly driven by dispersal events, in addition to host switching between plant families. We assume multiple colonisation events in the Middle Miocene, suggesting a particularly dynamic biogeographical history of *Melanastera* during this period. The colonisation of the Atlantic Forest was particularly important for the evolution of *Melanastera*. Our diversification rate analyses revealed a unique rate shift that may have occurred either at the time of *Melanastera* origin or during a major colonisation of the Atlantic Forest at 17.2 Ma in the clade ‘*Melanastera* crown group’. This colonisation coincides with a major host shift from *Melanastera* to the Melastomataceae family (Fig. 2), and together these events may have created new ecological opportunities for the diversification of *Melanastera*.

The next major colonisation event may have taken place from the Atlantic Forest back to the Amazon Forest (group MIV). During this relatively short time span, *Melanastera* underwent a high number of dispersal events, suggesting that the Atlantic and Amazon Forests played a fundamental role in the evolution of the genus. The colonisation events from these areas probably facilitated the diversification of *Melanastera*, which is reflected in the species-rich part of the tree. Thus, almost two-thirds of the *Melanastera* species analysed are endemic to the Atlantic and Amazon Forests suggesting that these areas have contributed to an accumulation of species diversity over time. The host shifts of *Melanastera* from the family Melastomataceae to Annonaceae (groups MV and MVI) in the Middle Miocene (Fig. 2) overlap with the colonisation of the Amazon Forest. This transition is also supported by the higher species richness of Annonaceae in the Amazon Forest, compared to the Atlantic Forest or the Cerrado biome (Ferreira et al., 2024; “Flora do Brasil,” 2020).

One possible explanation for the multiple dispersal events in *Melanastera* could result from its ability to spread over long distances. Although psyllids have never been suggested as good fliers, they are known to be passive dispersers that can perform long-distance movements with the wind (Percy, 2018, 2003). The experimental studies on the dispersal capacities of Asian Citrus and the African Citrus psyllids, *Diaphorina citri* (Psyllidae) and *Trioza erytreae* (Triozidae), respectively, showed that these species are able to disperse over more than 2 km (Antolinez et al., 2021; Nunes et al., 2023). In other insect groups, which are also known to be poor fliers, dispersal abilities have been demonstrated even more impressively. For example, it has been shown that different species of *Anopheles* mosquitoes (Diptera: Culicidae) can reach distances of up to 300 km (Huestis et al., 2019) and therefore such possibilities cannot be ruled out for psyllids.

The biogeographical analysis indicates that some regions of the Neotropics were recolonised several times (Fig. 5). In particular, the Atlantic Forest includes the highest number of *Melanastera* representatives (30 spp.) and was colonised at least five times independently. It is possible that the highest number of species in this area is the result of intensive sampling and that therefore the addition of more representatives from other areas to the analysis could increase the number of colonisation events also to other areas. Although part of the *Melanastera* diversity in the Atlantic Forest can be explained by the processes of adaptive radiation, the endemic diversity in this area, at least from the end of the Upper Miocene to the Pliocene (∼8–4 Ma), is probably due to a low and constant diversification that rather serves as a museum of diversity for *Melanastera*. This pattern is reflected in the presence of clades of closely related species in a single area or in neighbouring areas. The possible mechanisms that led to the local diversification of these clades can be explained by the Quaternary glaciation refugia hypothesis, according to which glaciation cycles in the Pleistocene speciation favoured the fragmentation of ecosystems and thus the speciation of some clades in allopatry (Garzón-Orduña et al., 2014; Toussaint et al., 2019; Wilson et al., 2012).

## 5. Conclusions

In the current study, we have produced a comprehensive, time-calibrated molecular phylogeny of the tropical and subtropical, species-rich psyllid genus *Melanastera* (Liviidae), which belongs to a taxonomically difficult and little-studied group of plant-sucking insects, the psyllids. Our results suggest that *Melanastera* likely originated at the beginning of the Miocene or at the Middle Oligocene in the Pacific region of the Neotropics, although most of its species diversification took place in the Upper Miocene.

Overall, the diversification and biogeographic history of *Melanastera* is broadly consistent with diversity scenarios proposed for other Neotropical clades (Antonelli et al., 2018; Condamine et al., 2015; Matos-Maraví et al., 2021b; McKenna and Farrell, 2006). *Melanastera* underwent an initial rapid burst of diversification either at its evolutionary origin or with its major shifts to new host families (Melastomataceae and Annonaceae) and colonisation of new adaptive zones (Atlantic and Amazon Forests), followed by a subsequent slowdown in speciation rates. We hypothesise that these shifts, which were accompanied by multiple dispersal events rather than vicariance, may have led to an accelerated radiation of the group in the Neotropics. Furthermore, our results suggest that local diversification in the Atlantic and Amazon Forests on closely related hosts from the species-rich families Annonaceae and especially Melastomataceae may have contributed to an accumulation of species diversity over time, ultimately leading to the current species richness in *Melanastera*.

## Authorship contribution statement

**Liliya Štarhová Serbina:** conceptualization, methodology, formal analysis, investigation, resources, data curation, writing – original draft, visualization, funding acquisition. **Daniel Burckhardt:** conceptualization, investigation, resources, writing – review and editing, supervision. **Lenka Petráková Dušátková:** methodology, investigation. **Dalva L. Queiroz:** investigation, resources. **Renato Goldenberg:** investigation, writing – review and editing. **Hannes Schuler:** resources, writing – review and editing. **Diana M. Percy:** investigation, resources, writing – review and editing. **Igor Malenovský:** conceptualization, investigation, resources, writing – review and editing, supervision.

## Acknowledgement

We are very grateful to M.L. Brotto and J.T.W. Motta (Museu Botânico Municipal, Curitiba, PR) for the identification of our Brazilian plant specimens. We gratefully acknowledge receiving the following collecting authorisations in Brazil: CNPq, IBAMA / SISBIO (11832; 37053). We thank Kristína Křížová and Kristýna Koukalová for their great help in organising the laboratory work and molecular sequencing at Masaryk University. LŠS was partially funded by a grant from the Swiss National Science Foundation (SNSF; P2BSP3_168733).

## Supplementary files

**Table S1.** List of the analysed psyllid species with the information on the host plant family and species and the distribution with federal states or provinces in the parentheses. The host plants for which the presence of immatures was confirmed are marked with an asterisk (*). The GenBank accession numbers in bold correspond to the gene fragments that were sequenced in the current study.

**Table S2.** Primers and PCR conditions.

**Table S3.** BioGeoBEARS dispersal multiplier matrices used in this study. A, Paraná region; B, Chacoan region; C, Boreal Brazilian region; D, South American transition zone; E, Mesoamerican region; F, Palaearctic region; G, Pacific region; H, Southeastern Amazonian region, I, Californian region.

**Table S4.** Characteristics of the multi-locus dataset.

**Table S5.** Summary of the divergence time estimates. ML = maximum likelihood analysis; BI = Bayesian inference. Upper and lower bounds of the 95% highest posterior density (HPD) in parentheses.

**Fig. S1.** Time-calibrated phylogenetic tree of the genus *Melanastera* with outgroups, based on Bayesian inference from molecular characters, with median ages and 95% credibility intervals, as inferred under a Yule model. Main clades are indicated in black. Fossil calibration points indicated by grey dashed lines.

**Fig. S2.** Credible set of macroevolutionary rate shift configurations with a prior of 1 shift (see Table 1).

**Fig. S3.** Spectral density plot of *Melanastera* and corresponding eigenvalues ranked in descending order. There is a gap between the seven and eight eigenvalues (indicated by an arrow), suggesting seven branching patterns (or modalities) in the phylogeny.

**Fig. S4.** Best phylogenetic tree of Melastomataceae, modified from Penneys et al. (2022). The tribes with genera of Melastomataceae associated with *Melanastera* representatives are indicated with blue arrows.

