## Supplementary figures and images for "Deciphering the patterns and timing of diversification of the genus *Melanastera* (Hemiptera: Psylloidea: Liviidae) in the Neotropics"

### Supplemental figure S1

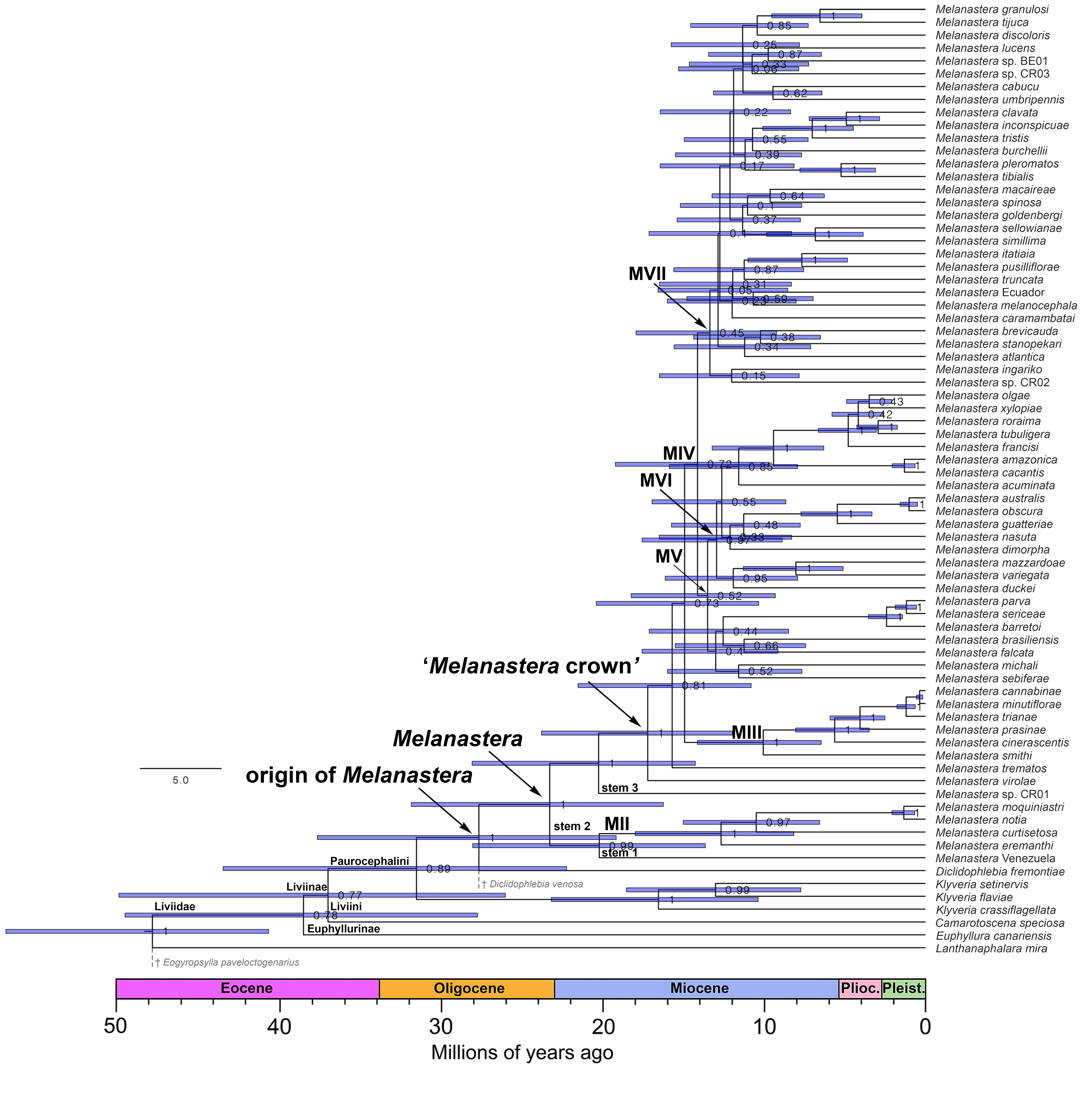

### Supplemental figure S2

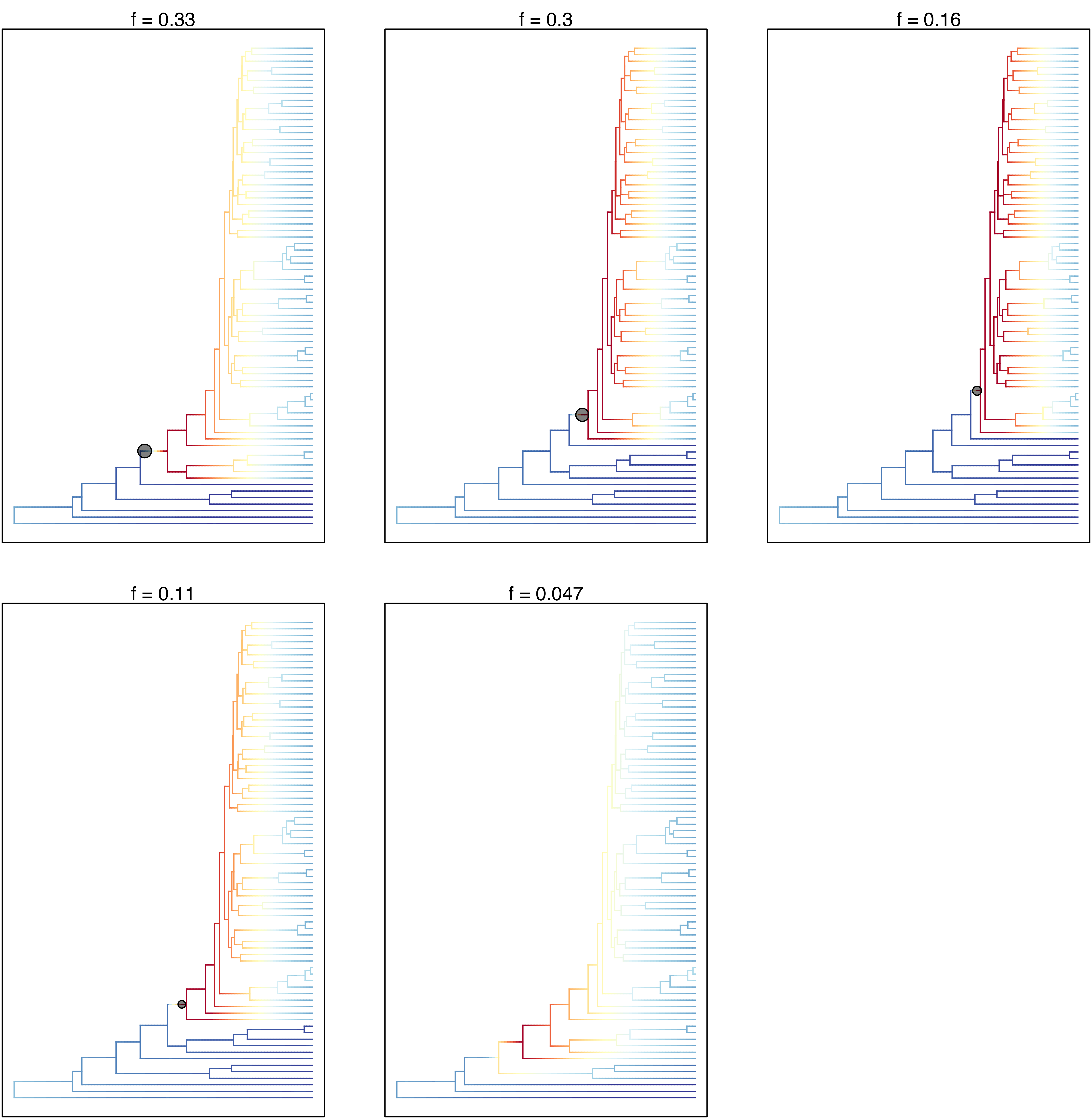

### Supplemental figure S3

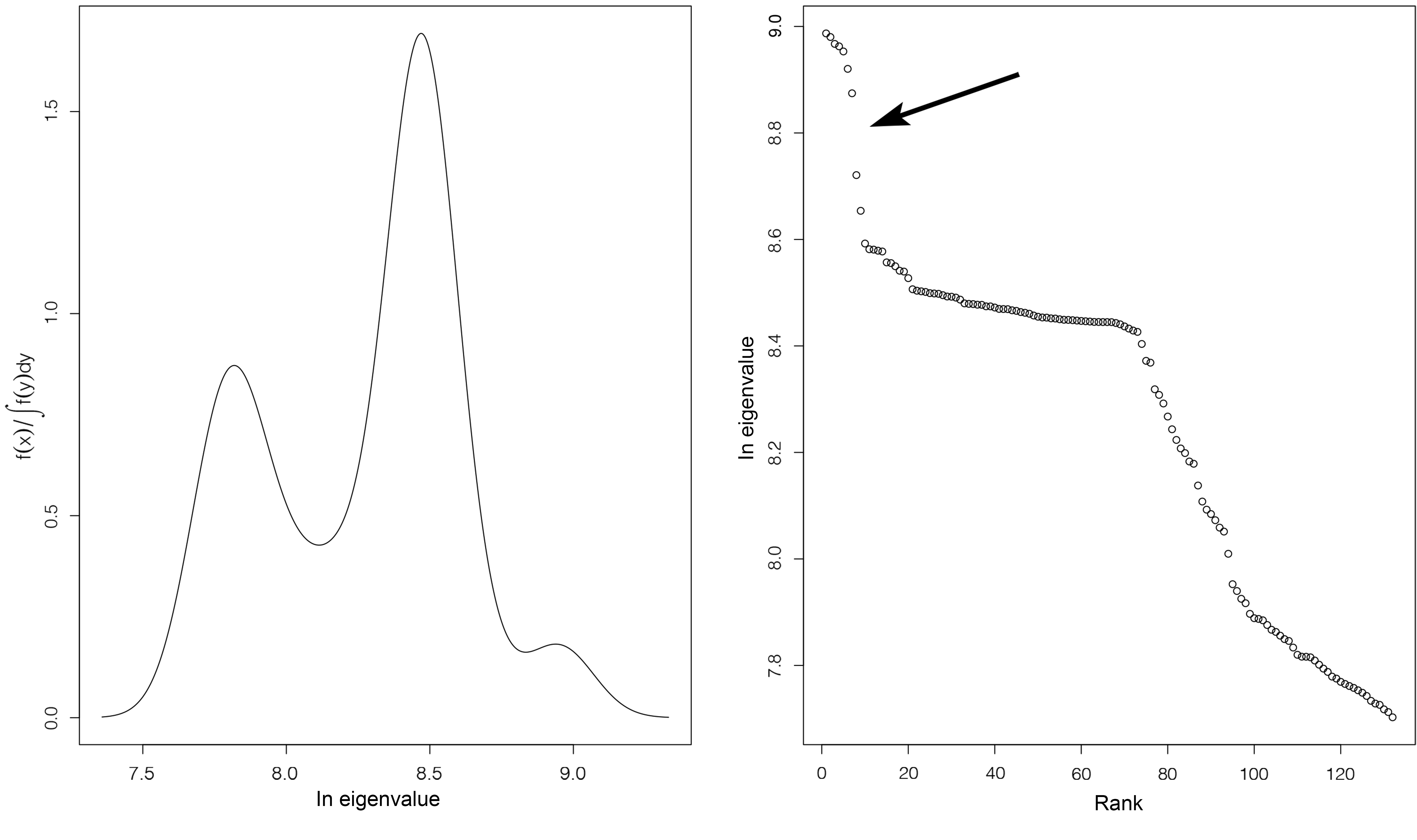

### Supplemental figure S4

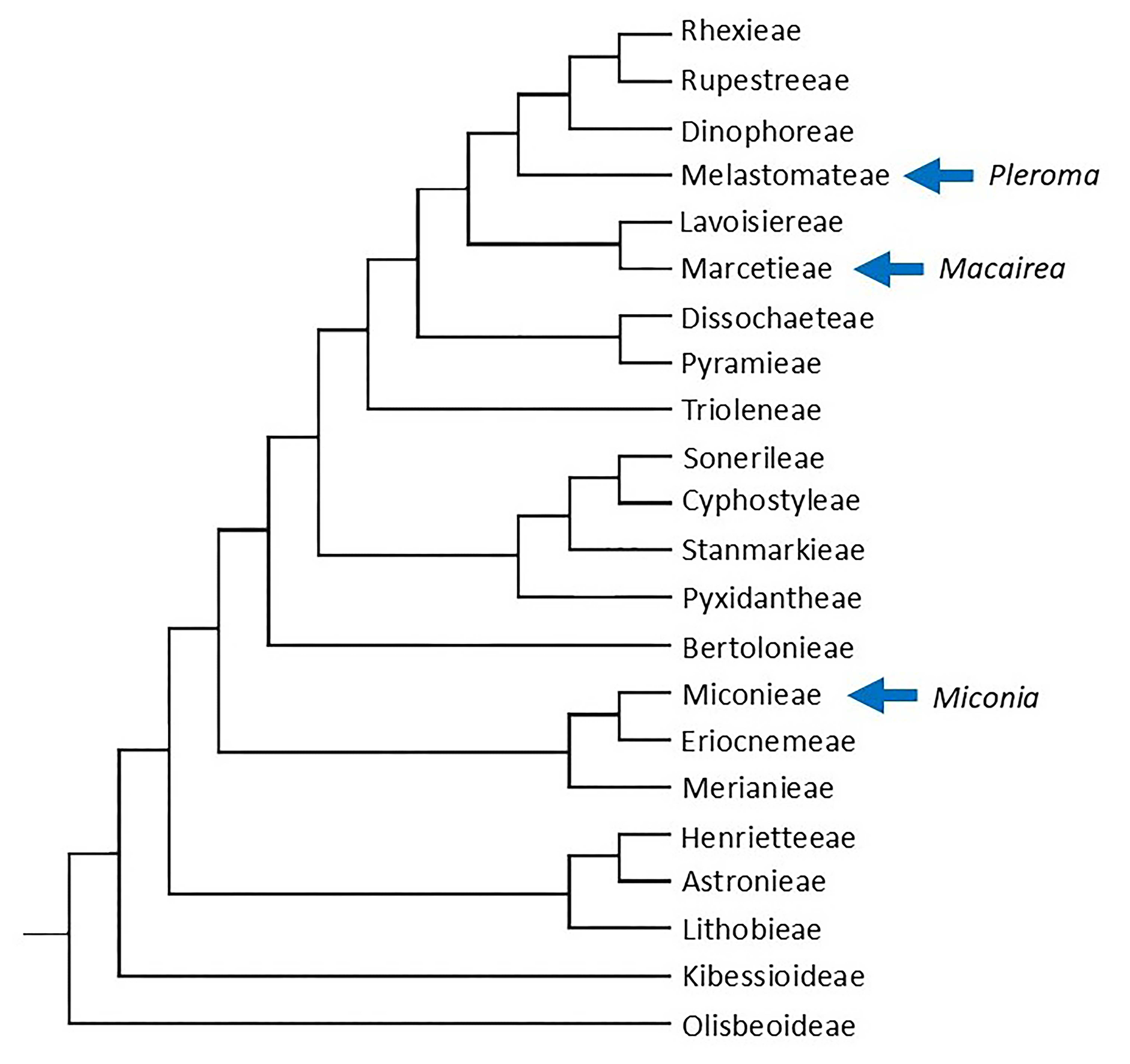
